# Human AUTS2 regulates neurodevelopmental pathways via dual DNA/RNA binding

**DOI:** 10.1101/2025.10.07.681056

**Authors:** Veronica Aparecida Monteiro Saia Cereda, Amandeep Sharma, Keegan Flanagan, Sara Elmsaouri, Sheila Steiner, Simone Benassi, Guilherme Reis-de-Oliveira, Jack T. Whiteley, Amoolya Chandrabhatta, Ana Paula Diniz Mendes, Lynne Randolph-Moore, Dimitra Xenitopoulos, Milla Jetzer, Ruth Oefner, Christopher Benner, Renata Santos, Gene W Yeo, Maria Carolina Marchetto

## Abstract

The *AUTS2* gene is implicated in neurodevelopmental and psychiatric disorders, with patient mutations leading to intellectual disability, microcephaly, and autistic behavior. While AUTS2’s chromatin-and RNA-related functions are recognized, its direct binding to RNA in human neural progenitors has not been previously demonstrated. Here, we used ChIP-seq and eCLIP-seq in human neural progenitor cells (NPCs) to map AUTS2’s chromatin targets and, for the first time, its direct RNA interactome. AUTS2 knockdown in NPCs led to widespread gene expression changes and impaired cell proliferation, migration, and neurite outgrowth. Integrated analysis revealed downregulation of Wnt pathway genes, notably *WNT7A*, among targets directly bound by AUTS2 at both chromatin and RNA levels. Supplementation with WNT7A rescued cellular phenotypes in AUTS2-deficient NPCs, underscoring the significance of Wnt signaling. These findings highlight AUTS2’s central role in human neurodevelopment and provide mechanistic insight into how its disruption may contribute to the pathology of neurodevelopmental disorders.

## Introduction

The *AUTS2* (Autism Susceptibility Candidate 2) gene was first identified as a risk factor for autism spectrum disorder (ASD) and has since been implicated in brain development, a variety of neurodevelopmental and psychiatric disorders, and mammalian evolution ^1–4^. Genomic structural variants and single-nucleotide polymorphisms (SNPs) in this gene cause AUTS2 syndrome, a dominantly inherited neurodevelopmental disorder marked by symptoms including developmental delay, intellectual disability, microcephaly, feeding difficulties, and craniofacial abnormalities ^1,5^. The human *AUTS2* gene encodes two primary isoforms: a full-length isoform derived from exons 1–19 and a shorter C-terminal isoform generated from an alternative transcription start site in exon 9. Mutations in the 3’ region of *AUTS2* (exons 9–19) are associated with greater disease severity ^6,7^. In mice, deletions affecting only the long isoform result in relatively mild developmental phenotypes, whereas mutations affecting the short isoform lead to more severe brain pathology, including cognitive, memory, and social deficits ^8–10^. Additionally, neurodevelopmental defects have been documented in zebrafish with knockdown of *auts2* ^11^. Although these clinical and experimental findings implicate AUTS2 in diverse aspects of brain development, studies dissecting its cellular and molecular functions remain limited in number and are largely centered on mouse models. Understanding AUTS2’s mechanism is further complicated by the existence of distinct isoforms and cellular localizations (nuclear and cytoplasmic), depending on cell type and developmental stage ^10,12–14^.

AUTS2 has been described to act as a transcriptional activator or repressor depending on the cellular context ^2,8,10,12,13,15^. Studies in mouse brain, *in vitro* cellular systems (such as HEK293 and mouse embryonic stem cells), and yeast two-hybrid assays showed that AUTS2 is recruited to promoter regions of actively transcribed genes, where its interaction with P300, CK2, and NRF1 proteins within the non-canonical Polycomb repressor complex 1 (PRC1.3/1.5) core activates transcription ^2,8,15,16^. Genes targeted by AUTS2-PRC1 are significantly enriched for neurodevelopmental functions ^2,8^. Recently, evidence has shown that AUTS2 interacts with the Polycomb complex PRC2 in mouse cortical intermediate progenitors, mediating chromatin modification and transcriptional repression, which are necessary for proliferation and upper-layer neuron specification ^14^. AUTS2 is also found in RNA-protein complexes and binds transcripts involved in RNA metabolism required for cortex and dentate gyrus development ^9^. Whether AUTS2 binds RNA directly or regulates the activity of associated complexes is still unclear. In the cytoplasm, AUTS2 modulates actin dynamics and neuronal migration by regulating Rho-family GTPases, notably activating Rac1 for neurite extension and cortical lamination ^10^. Also, AUTS2 restricts excitatory synapse formation in forebrain pyramidal neurons, shaping dendritic spine density and ensuring proper balance of excitation/inhibition in neural circuits ^4^. Collectively, mouse studies establish AUTS2 as a multifunctional regulator coordinating transcription, epigenetics, RNA metabolism, actin dynamics, and synaptogenesis at critical phases of brain development, from neural progenitor differentiation and migration to neuronal maturation, circuit assembly, and cortical lamination.

In the human context, only two studies have investigated AUTS2-related neurodevelopmental defects using cerebral organoids, both revealing abnormalities during early brain development. One study modeled the pathogenic missense variant T534P, which causes microcephaly and severe intellectual disability, using patient-derived induced pluripotent stem cells (iPSCs) ^17^. The other study employed CRISPR/Cas9-mediated deletion of *AUTS2* in human embryonic stem cells (hESCs) to study its role in neuronal differentiation ^18^. Both studies found a reduction in organoid size, impaired neural progenitor populations, and dysregulation of WNT/β-catenin signaling; however, the underlying mechanisms seem to differ. The heterozygous missense variant caused a reduction in progenitor proliferation, promoted premature neuronal differentiation, and lowered WNT/β-catenin pathway activity, as revealed by single-cell RNA sequencing (scRNA-seq) ^17^. This mutation also caused a disruption in the interaction with P300, likely altering transcriptional activity of the AUTS2-PRC1 complex. In contrast, complete loss of AUTS2 resulted in a shift of neuroepithelial cell differentiation into NPCs to differentiation into choroid plexus-like cells through hyperactivated WNT/β-catenin signaling ^18^. These findings support a major role of AUTS2 in regulating developmental NPC functions and cortical neuronal differentiation, and the discrepancies highlight the complexity of AUTS2 function and the need for further studies.

Here, we present a comprehensive genome-wide analysis of AUTS2 function in human cortical NPCs derived from pluripotent stem cells. We show that human AUTS2 not only interacts with DNA but also binds directly to RNA, expanding its known regulatory mechanism beyond what has been described in mouse models. By generating AUTS2 knockdown cell lines, we identified differentially expressed genes that are targeted by AUTS2 at both DNA and RNA levels, revealing a network of neurodevelopmental genes and Wnt signaling components impacted by AUTS2 loss of function. Among these, *WNT7A* emerged as one of the most strongly repressed genes following AUTS2 knockdown. Supplementation with recombinant WNT7A rescued the phenotypes caused by AUTS2 deficiency, confirming its role in AUTS2-derived pathologies. Together, these findings add evidence to establish AUTS2 as a central coordinator of neurodevelopmental gene networks in human NPCs, with regulation of the Wnt pathway as a critical mechanism underlying its role in cortical development.

## Results

### AUTS2 associates with active chromatin in human neural progenitors

Chromatin immunoprecipitation followed by genome-wide analysis (ChIP-seq) revealed 702 high-confidence AUTS2 binding sites, with the majority (36%) located within promoter regions proximal to transcription start sites (TSS) in NPCs derived from hESC H9 line (**Figs. 1A, B,** and **Supplementary Table 1)**. ChIP profiles of chromatin modifications associated with AUTS2 peaks revealed colocalization with accessible chromatin (ATAC-seq), RNA polymerase II and histone modifications associated with active transcription, including histone H3 acetylated at lysine-27 (H3K27ac), and histone H3 trimethylated at lysine-4 (H3K4me3) (**Fig. 1C**, **Supplementary Fig. 1A**). Importantly, there was minimal association between AUTS2 peaks and histone H3 lysine-27 trimethylation (H3K27me3), an epigenetic mark associated with gene repression and heterochromatin (**Fig. 1C**, **Supplementary Fig. 1B**). Further investigation of the ChIP-seq data using Hypergeometric Optimization of Motif EnRichment (HOMER)^19^ indicated that AUTS2 preferentially binds near active regulatory elements that are strongly enriched for DNA motifs recognized by the RFX, SOX and NR4A1 families of transcription factors, which have critical functions in neuronal development (**Fig. 1D**). AUTS2 binding sites were also enriched near genes critical for neurodevelopmental processes, including axonogenesis, forebrain development, and neuronal differentiation (padj-value < 0.05) (**Fig. 1E**). Examples of AUTS2 target genes identified in human NPCs included: *ASCL1, BCL11B, GATA2*, *HES5*, *NEUROG1*, *NEUROD1*, *NFIB, NKX2-2, NOTCH2*, *OTX2*, *ROBO2, SOX1*, *WNT3A* and *WNT7A* (**Fig. 1F**). In conclusion, AUTS2 in human NPCs predominantly binds to regions near TSSs of genes that are essential for neurodevelopment. These sites are enriched in active chromatin marks, supporting a role for AUTS2 in positively regulating the transcriptional programs of the developing central nervous system.

**Fig. 1:**
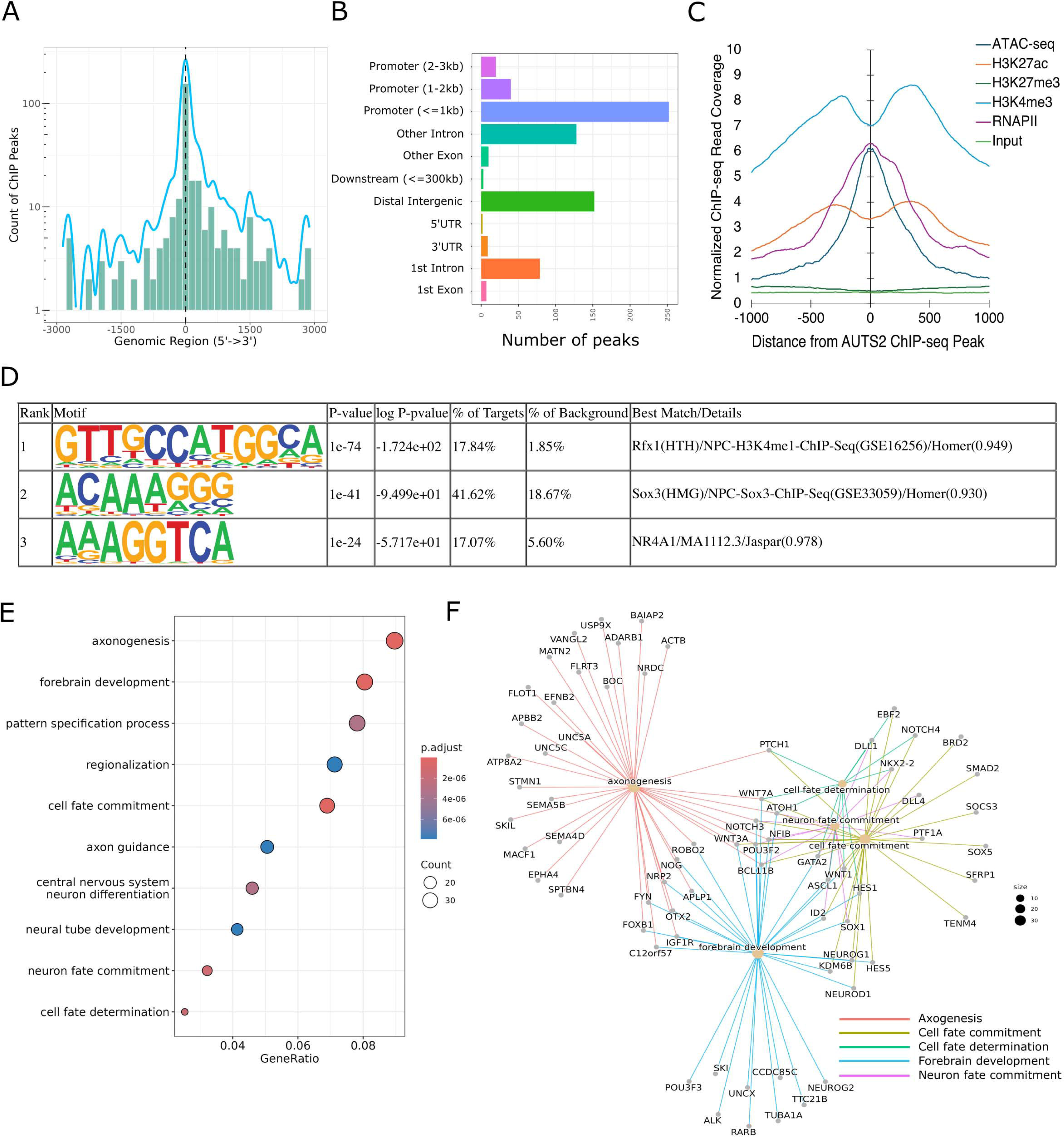
AUTS2 ChIP-Seq analysis in human NPCs revealed enrichment of proteins associated with neurodevelopmental processes. **A)** Distribution of ChIP peaks as a function of distance from the TSS. Negative values represent peaks upstream of the TSS (position 0 in the x axis). **B)** Barplot displaying feature type counts for AUTS2 ChIP targets. **C)** ChIP reads density plots (y axis) for levels of indicated histone marks and RNA polymerase II (RNA PII) at loci targeted by AUTS2. **D)** Top *de novo* enriched DNA motifs in AUTS2 peaks analyzed by HOMER. Motif enrichment and associated p-values are calculated using the cumulative binomial distribution. **E)** Dotplot showing GO pathway enrichment of ChIP target genes. The color of the dots represents the significance of enrichment, while size represents the number of targets present in the pathway. P-adjust were calculated using an over-representation test and the Benjamini–Hochberg correction, implemented in clusterProfiler 4.12.6^53^. **F)** Network diagram showing target genes in the top 5 significantly enriched GO pathways.

### AUTS2 binds to transcribed RNA from neurodevelopmental genes

AUTS2 was previously implicated in RNA metabolism in neonatal mouse cortex but the nature of this interaction has remained unclear, as RNA immunoprecipitation followed by high-throughput sequencing (RIP-seq) analyses could not reliably distinguish between direct and indirect RNA binding ^9^. To overcome this limitation and explore the relevance of AUTS2’s RNA interactions in the human context, we performed enhanced cross-linking immunoprecipitation followed by high-throughput sequencing (eCLIP-seq) ^20^ in human NPCs. eCLIP-seq offers a robust and standardized framework for generating high-resolution, genome-wide maps of RNA-binding protein (RBP) interaction sites. Using this technique, we were able to comprehensively map AUTS2’s RNA interactome, which represents the first in depth characterization of human AUTS2 as an RBP. AUTS2 eCLIP-seq analysis identified 883 binding sites (**Fig. 2, Supplementary Table 2**). Metagene analysis suggests that AUTS2 preferentially binds near the TSS or transcription termination site (TTS) of its target RNAs (**Fig. 2A**). This pattern is consistent with co-transcriptional deposition, where there is simultaneous or coupled processing of both DNA transcription and the addition or modification of molecules or factors to the newly synthesized RNA molecule at the TSS and TTS^21^. In addition, identification of AUTS2-binding sites within introns (**Fig. 2B**) suggest a potential role for AUTS2 in post-transcriptional regulatory processes, such as pre-mRNA splicing or nuclear RNA processing. Enriched RNA-bound sequences by AUTS2 revealed an enrichment of gene transcripts known to be important for neurodevelopmental processes, including neurogenesis, axonogenesis, forebrain development, and neural precursor cell proliferation (**Fig. 2C**), similar to pathways identified in the ChIP-seq analysis. Examples of gene transcripts bound by AUTS2 in human NPCs include *HES6*, *SOX11*, *FOXP2*, *CTNNB1*, *CUX1*, *NOTCH3*, *VIM*, *CDH2*, *NOVA2*, *WNT7A* and *AUTS2* itself; most of which are implicated in critical neurodevelopmental processes and disabilities (**Fig. 2D**). Collectively, these analyses demonstrate that the human AUTS2 possesses both DNA and RNA binding capabilities and can be characterized as a DNA-and RNA-binding protein (DRBP). This definition expands AUTS2’s previously described functions and suggests a dual regulatory role, potentially influencing the expression of multiple genes involved in human neurodevelopment at multiple levels.

**Fig. 2:**
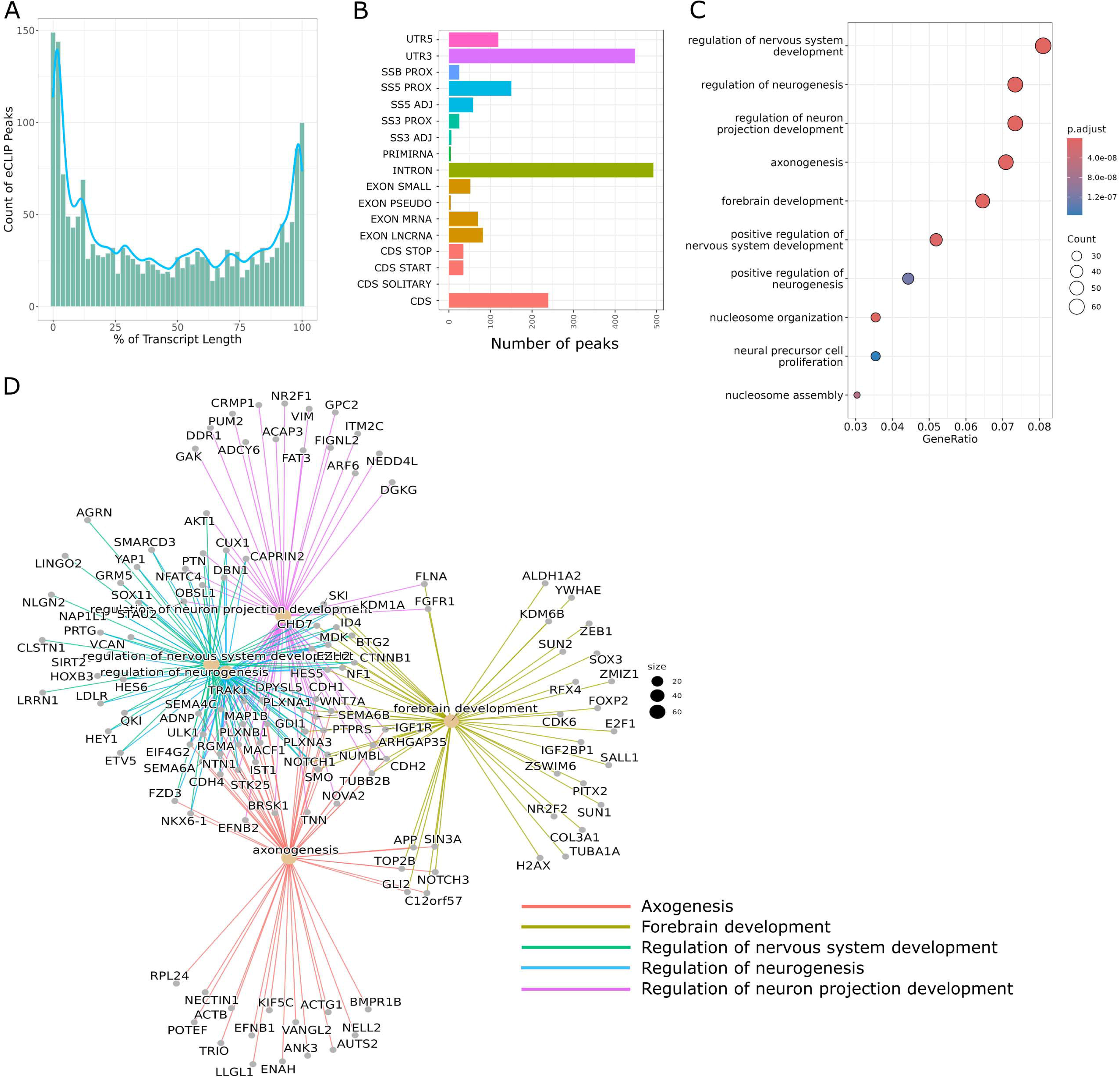
eCLIP analysis revealed AUTS2 as an RNA-binding protein in human NPCs. **A)** Histogram displaying the distribution of eCLIP peaks as a function of transcript length. Binning is based on percentage (%) of transcript length, with 0 being the TSS and 100 being the TTS. **B)** Barplot displaying feature type counts for AUTS2 eCLIP targets. **C)** Dotplot showing GO pathway enrichment of eCLIP target genes. The color of the dots represents the significance of enrichment, while size represents the number of targets present in the pathway. p-adjusted are calculated using a gene-ontology over representation test (Fisher’s exact test) and adjusted using the Benjamini-Hochberg method. **D)** Network diagram showing target genes in the top 5 most significantly enriched GO pathways.

### Loss of AUTS2 disrupts neuronal differentiation pathways in human neural progenitors

To investigate the impact of the functional role of the AUTS2 expression protein in human neurodevelopment, we performed CRISPR/Cas9 genome editing in hESCs using guide RNAs (gRNAs) targeting *AUTS2* gene as well as non-targeting gRNA to generate control clonal lines. Guide RNAs specifically targeted exon 9 and exon 12 of the *AUTS2* gene given the established link between mutations in the 3’ region of AUTS2 (exons 9–19) and heightened disease severity, coupled with their predicted importance in neuronal differentiation^6,7^.

Following Cas9-mediated cleavage and successful homologous recombination repair, hESC clones containing targeted insertions/deletions (INDELs) within the *AUTS2* gene were isolated and expanded (experimental design illustrated on **Fig. 3A**). We next employed the Amplicon-EZ assay for targeted next-generation sequencing (NGS) to screen for successful edits efficiently. Out of all the clones analyzed, three showed the biallelic modifications in the *AUTS2* gene: one clone with an INDEL in exon 9 and two clones harboring INDELs within exon 12 in both alleles (**Supplementary Fig. 2**). Western blotting showed a significant decrease in AUTS2 protein expression in all three edited clones compared to control lines (**Fig. 3B**). These results indicate that the introduced INDELs effectively blocked AUTS2 expression, providing a model system to study the role of human AUTS2 in neurodevelopment and disease. Western blot analysis in cortical NPCs revealed a protein band at approximately 79 kDa, consistent with the short AUTS2 C-terminal isoform previously characterized during embryonic brain development in mice ^6,10^. We confirmed the presence of the AUTS2 C-terminal isoform with a TSS in exon 9 using 5’-RACE (**Fig. 3C and Supplementary Fig. 3**). These findings are consistent with previous reports by Beunders et al. (2013) and Biel et al. (2022)^6,22^ showing a transcribed region spanning exons 9 to 19 encompassing 2136 base pairs, encoding a 711 amino acids protein.

**Fig. 3:**
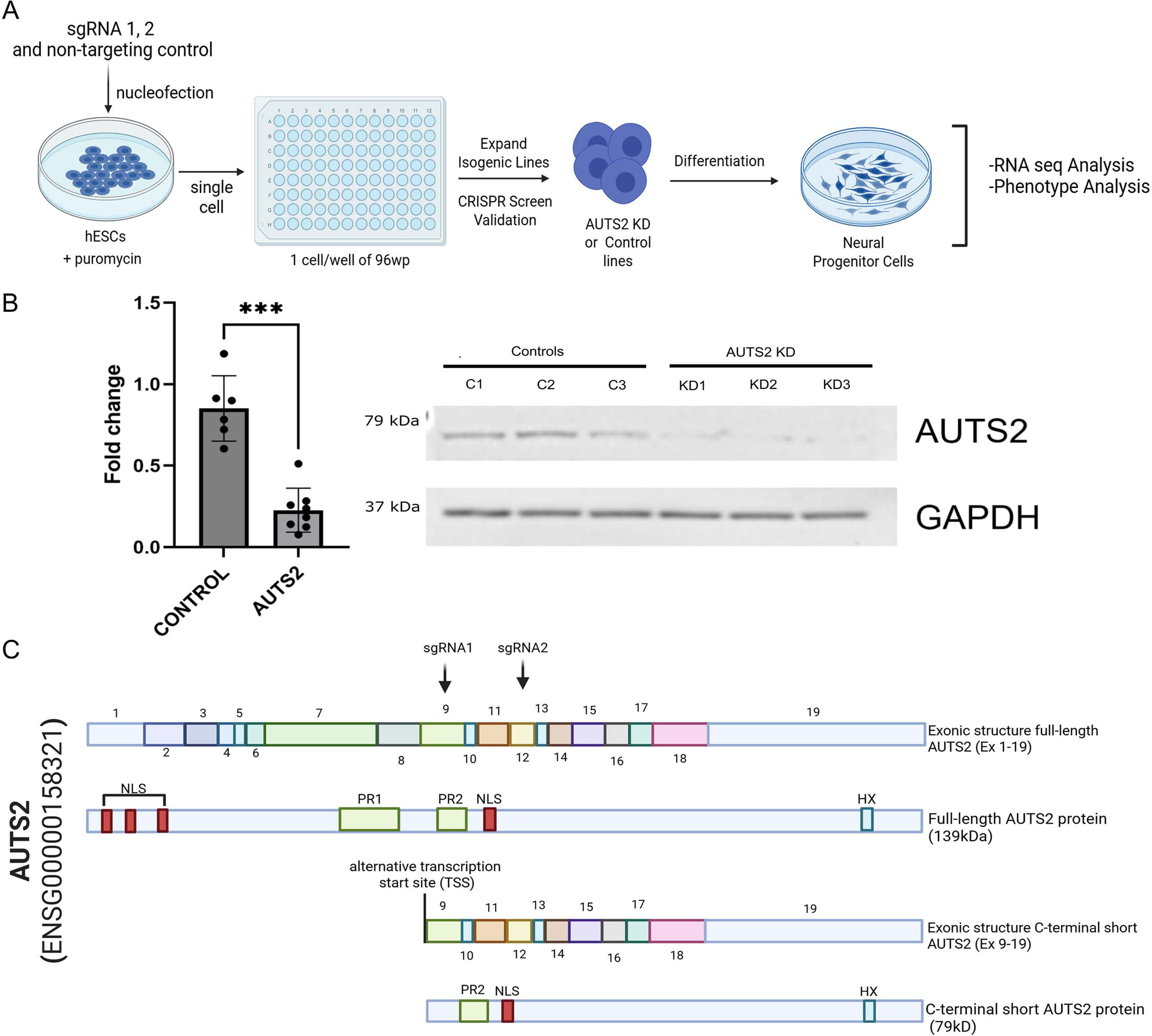
CRISPR-Cas9 targeting of *AUTS2* gene to generate homozygous mutants with reduced mRNA and protein. **A)** Schematic of CRISPR-Cas9 experimental design. This panel outlines the step-by-step process used to generate AUTS2 knockdown (KD) and non-targeted control lines. **B)** Western blotting analysis of AUTS2 protein in NPCs generated from three control lines and three AUTS2-KD (knockdown) lines. p-values were determined by unpaired t-test with Welch’s correction (n=3 cell lines, 3 independent experiments). *p<0.05; **p<0.01; ***p<0.001; ****p<0.0001. **C)** Schematic depiction of the exonic structure of 2 distinct AUTS2 transcript variants and its respective protein products. The 2 top sequences represent full length transcript (exons 1-19) and full length protein (139 kDa). The bottom sequences represent the shorter variant of AUTS2 (alternative TSS, exon 9-19) and its respective shorter C-terminal protein isoform (79kDa). This schematic shows key functional domains within these sequences, such as the alternative TSS, nuclear localization signal (NLS), which directs the protein to the nucleus, and proline-rich (PR) domains that may mediate protein-protein interactions. Panels **A** and **C** were created using <u>BioRender.com</u>.

RNA sequencing (RNA-seq) analysis in NPCs comparing control samples to those with AUTS2 knockdown identified 5,375 differentially expressed genes with statistically significant changes (pAdj-value < 0.05) and a fold change greater than 2. Out of the differentially expressed genes, 3,558 genes were upregulated, while 1,817 genes were downregulated (**Fig. 4A and Supplementary Table 3**). We validated seven of the topmost differentially regulated genes by qPCR confirming the RNA-seq results (**Fig. 4B**). Pathway analysis of the differentially expressed genes revealed significant dysregulation in neurodevelopmental processes, including regulation of nervous system development, axonogenesis and axon development, and regulation of neurogenesis (**Fig. 4C)**. These pathways comprise a network of interacting genes (**Fig. 4D**), including selected validated genes: *WNT7A*, *HES5*, *PRTG*, *CFI*, *CRYBA1*, and *TRIM48*. Further analysis of the pathways affected by AUTS2 deficiency showed that genes within the gene ontology (GO) terms ‘cell cycle’, ‘RNA metabolism’ and ‘DNA methylation’, were generally downregulated, confirming a role for AUTS2 in neurogenesis, RNA metabolism, and epigenetics (**Figs. 4E, F**). In summary, transcriptomic analysis of AUTS2 knockdown in human NPCs reveals disruptions in canonical pathways involved in neuronal differentiation and RNA metabolism.

**Fig. 4:**
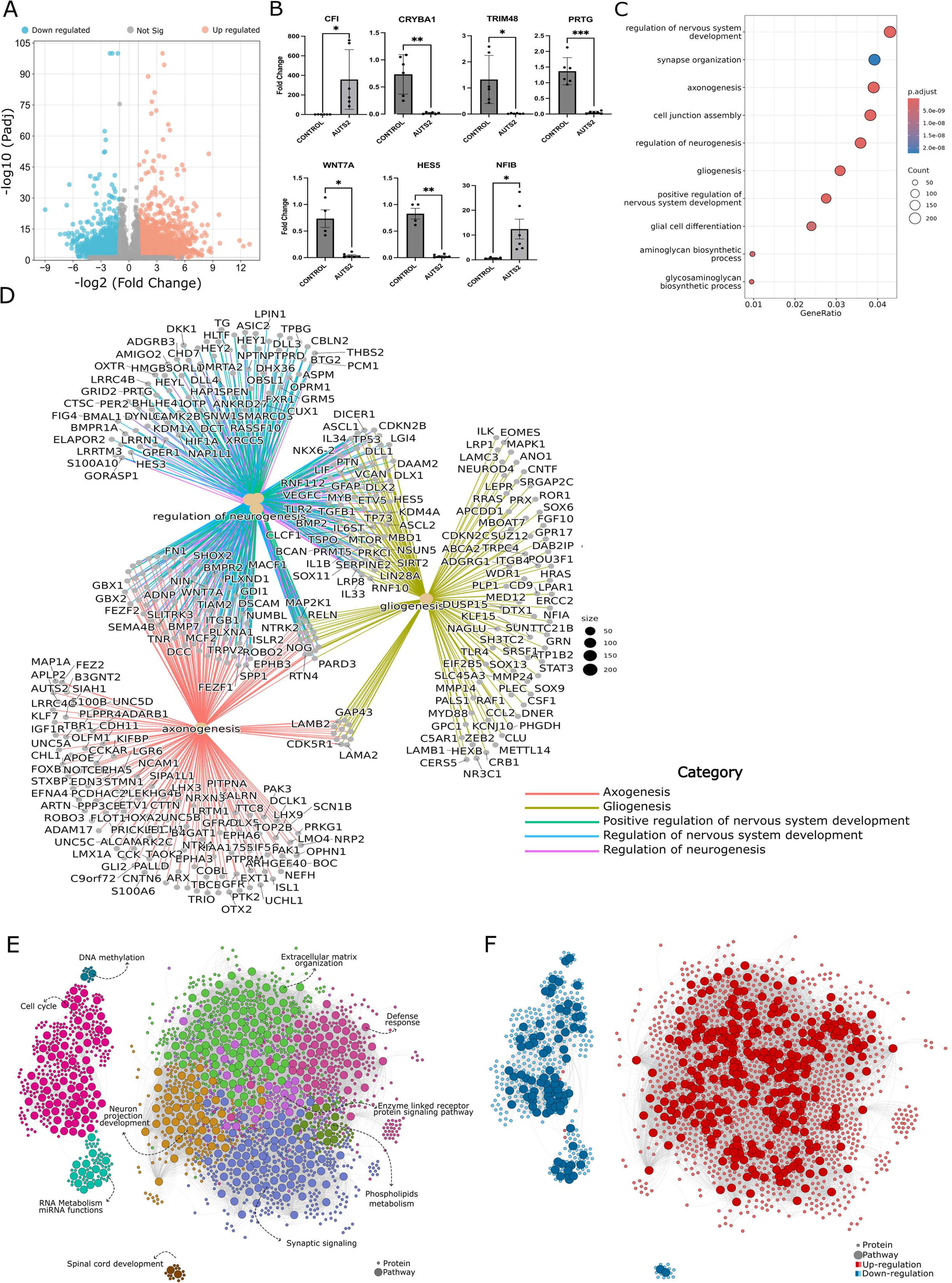
Transcriptomic analysis of AUTS2-deficient human NPCs showed deregulations in pathways related to neuronal differentiation. **A)** Volcano plot displaying differentially expressed genes in AUTS2-deficient versus control NPCs. Genes with an absolute log-fold change greater than 2 (i.e., mRNA with a fold change >2 or <−2) and padj< 0.05 are included. **B)** RT-qPCR analysis of mRNA expression of several top 20 genes found significantly differentially expressed in RNA-seq analysis. p values were determined by an unpaired t-test with Welch’s correction (n=3 cell lines in duplicate). *p<0.05; **p<0.01; ***p<0.001; ****p<0.0001. **C)** Dotplot showing GO pathway enrichment of differentially expressed genes. The color of the dots represents the significance of enrichment, while the size represents the number of targets in the pathway. p-adjusted are calculated using a gene ontology over-representation test (Fisher’s exact test) and adjusted using the Benjamini-Hochberg method. **D)** Network diagram showing target genes in the top 5 most significantly enriched GO pathways. **E)** Network visualization of GO pathway enrichment in RNA-seq data from AUTS2-deficient compared to control NPCs. Dot color indicates the enriched GO cluster, and dot size is proportional to the number of target genes within each pathway. **F)** Network representation of enriched pathways and genes, and their regulation, in RNA-seq data from AUTS2-deficient compared to control NPCs. Dot color represents gene expression changes (red for up-regulated and blue for down-regulated), while dot size is proportional to the number of target genes within each pathway.

### Dual DNA/RNA binding by AUTS2 regulates genes involved in neurodevelopmental disorders

To investigate AUTS2-mediated regulation at transcriptional and post-transcriptional levels, we integrated ChIP-seq, eCLIP-seq, and RNA-seq data to identify genes co-bound by AUTS2 at DNA and RNA levels whose expression is altered upon AUTS2 knockdown. A significant overlap was observed between the target genes identified by ChIP-seq and eCLIP-seq (**Fig. 5A**), with 69 genes targeted by AUTS2 both transcriptionally and post-transcriptionally. Analysis of the genomic distance between ChIP and eCLIP peaks for dual targets revealed a high degree of similarity in binding locations (**Fig. 5B**). We compared dual targets of AUTS2 with genes associated with neurodevelopmental disorders using the following data sets: 1) Simons Foundation Autism Research Initiative (SFARI) (p < 0.00965); 2) Deciphering Developmental Disorders (DDD) (p < 0.00531); and 3) the Intellectual Disability gene list within the ORPHANET database (ID) (p < 0.0213)^23–25^. There was significant enrichment for AUTS2 co-bound genes within all these lists (**Fig 5C, Supplementary Fig. 4A, B)**. We then calculated the enrichment score for a disorder that is not related to neurodevelopment (breast cancer, BC ^26^) and identified no association with AUTS2 co-bound genes (p < 0.831) (**Fig. 5C**). These analyses indicate that genes co-regulated by AUTS2 are involved in a broad spectrum of neurodevelopmental disorders. Of the 69 dual targets identified, 39 of them were also found to have significant differential expression following AUTS2 knockdown (**Fig. 5D and Supplementary Table 4**). Of the differentially regulated co-bound genes, over two-thirds (28 genes) were downregulated in AUTS2-deficient lines, supporting a potential role for AUTS2 in gene activation (**Fig. 5E**). Pathway analysis conducted using these 39 genes identified significant enrichment in pathways crucial for neurodevelopment, including Wnt signaling, neurogenesis, axonogenesis, and neuron migration (**Fig. 5F**). *WNT7A* was the most significantly differentially regulated gene in AUTS2 knockdown lines and is well established as a regulator of central nervous system development, influencing processes such as neuronal differentiation, axon guidance, and synapse formation^27^. We further investigated the *WNT7A* locus using the University of California Santa Cruz (UCSC) genome browser to visualize the sites of AUTS2 shared ChIP and eCLIP peaks (**Fig. 5G**). ChIP-seq peaks called by MACS2^28^ highlighted a broad enrichment at a *WNT7A* open-chromatin region overlapping with ATACseq and H3K27ac tracks, indicating that AUTS2 is interacting with a putative active enhancer region within the gene locus. eCLIP peaks called by Skipper^29^ revealed a sharp focal enrichment within the first intron of *WNT7A* transcript, and a pronounced H3K4me3 signal around the area reflects the peak’s proximity to the TSS (**Fig. 5G**). Investigation of other co-regulated gene loci (*i.e*. *NSD2*), show a similar pattern of ChIP-seq peaks overlapping with open chromatin (**Supplementary Fig. 4C**).

**Fig. 5:**
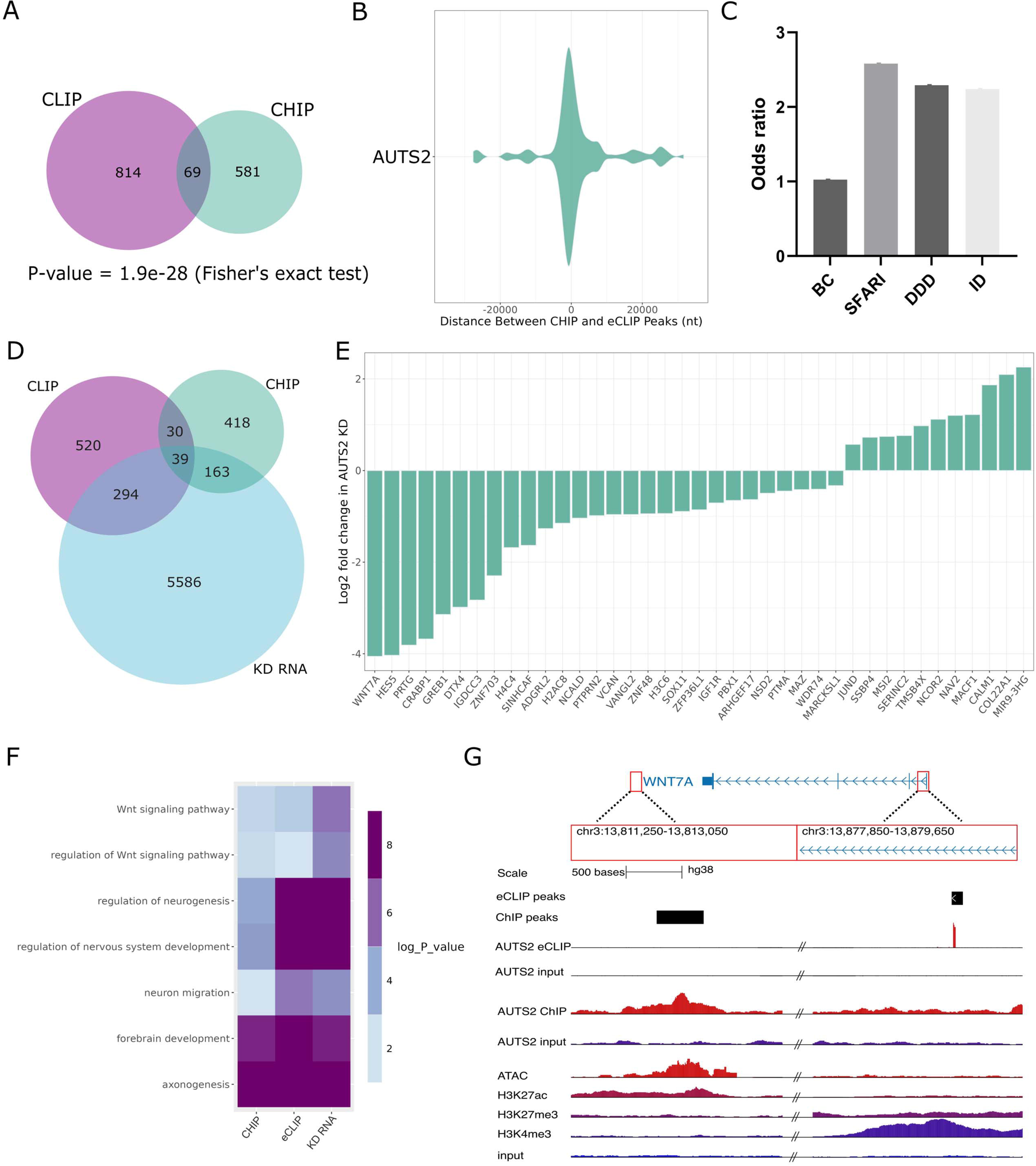
AUTS2 regulates gene expression at both transcriptional and post-transcriptional levels through its dual DNA-and RNA-binding activities. **A)** Venn diagram showing AUTS2 target genes identified via eCLIP or ChIP. Jaccard index of eCLIP and ChIP target overlaps, significance of eCLIP and ChIP overlap calculated via Fisher’s exact test. **B)** Distance between ChIP and eCLIP peaks within the same gene only. Distance is calculated as the difference in genomic position of ChIP versus eCLIP peaks (ChIP-eCLIP). Negative values represent eCLIP peaks downstream of ChIP peaks. **C)** Odds ratio of the overlap between genes related to developmental disorders and the AUTS2 interactome. The odds ratios represent the enrichment of genes associated with AUTS2 CLIP-seq/ChIP-seq experiments within three distinct gene datasets. The datasets include genes from the SFARI (Simons Foundation Autism Research Initiative), intellectual disability-related genes - ORPHANET (ID), and DDD (Deciphering Developmental Disorders) datasets, which represent developmental disorders, with breast cancer (BC) genes used as a negative control. **D)** Venn diagram showing AUTS2 eCLIP/ChIP target genes and genes with significant differential expression following AUTS2 knockdown. **E)** Barplot displaying log-transformed change in expression for 39 genes found to be dual eChIP/CLIP targets with significant differential expression following AUTS2 KD versus control NPCs. **F)** Heatmap showing gene-set enrichment for Wnt signaling and neurodevelopmental pathways across three gene sets (left to right): genes identified as significant eCLIP targets, significant ChIP targets, and genes differentially expressed following AUTS2 knockdown. p-values were calculated using a gene ontology over-representation test (Fisher’s exact test) and adjusted using the Benjamini-Hochberg method. **G)** UCSC Genome Browser view of the *WNT7A* locus (hg38) divided in two genomic windows (chr3:13,811,250–13,813,050 and chr3:13,877,850–13,879,650) for optimal visualization of both AUTS2-ChIP and-eCLIP peaks within the locus. The RefSeq annotation track shows exons as blue boxes and introns as lines. ChIP-seq peaks are called by MACS2 (see Methods) and appear as black boxes, with normalized coverage (IP in red, input in purple). eCLIP peaks called by Skipper (see Methods) are marked with black boxes, and normalized coverage (IP in red, input in purple). Also shown are ATAC-seq, H3K27ac, H3K27me3, and H3K4me3 tracks.

Together, these data reveal a novel role for human AUTS2 in regulating genes involved in neurodevelopment and disease at both the transcriptional and post-transcriptional levels through its dual DNA-and RNA-binding activities.

### AUTS2 deficiency causes impairment in cell proliferation, neuronal migration and neurite outgrowth

Our analyses strongly suggest that AUTS2 plays a central role in neurogenesis through coordinated DNA and RNA-level regulation of gene expression. To assess the cellular consequences of AUTS2 knockdown, we examined its impact on critical stages of neurogenesis: progenitor proliferation, migration, and neurite outgrowth. Cell cycle analysis revealed that AUTS2-deficient NPCs are arrested in the G1 phase, with a significant reduction in the S phase population compared to controls, indicating impaired cell cycle progression (**Fig. 6A**). Consistent with decreased cell proliferation and premature cell cycle exit, AUTS2 knockdown NPCs formed smaller neurospheres compared to the control group (**Fig. 6B**). Integration of ChIP, eCLIP, and RNA-seq datasets revealed significant enrichment of genes associated with cellular migration. To functionally validate this finding, we performed scratch assays on NPC cultures. AUTS2 knockdown NPCs exhibited delayed wound closure compared to controls, indicating impaired migratory capacity (**Fig. 6C**). Finally, we assessed neurite outgrowth and observed a significant reduction in neurite length in AUTS2-deficient neurons compared to controls (**Fig. 6D**), indicating impaired neuronal maturation and defective extension of neuronal projections. Together, these findings demonstrate that AUTS2 knockdown disrupts critical cellular processes underlying neurogenesis and neuronal differentiation.

**Fig. 6:**
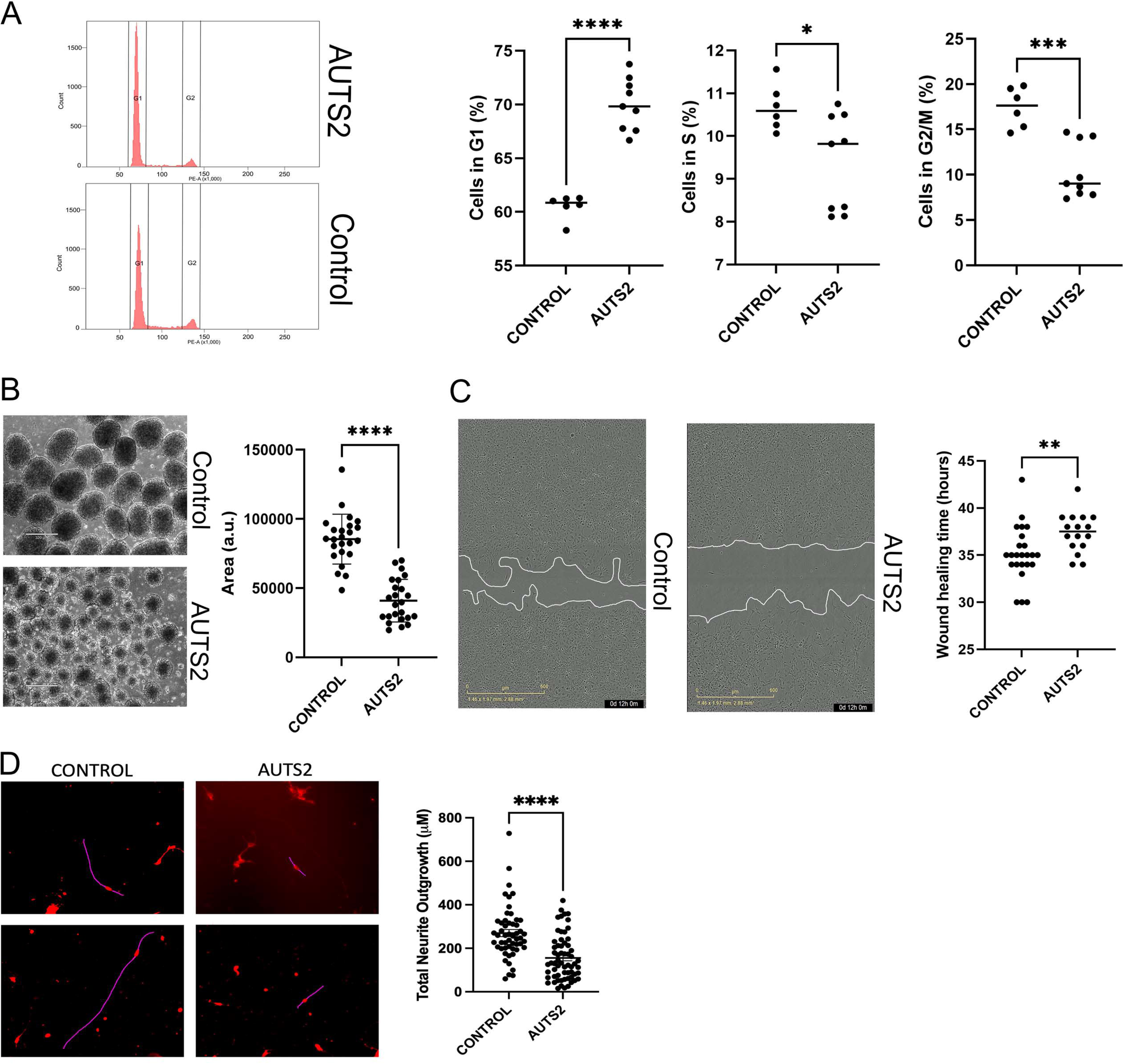
AUTS2 knockdown in NPCs results in deficits in proliferation, migration and neurite outgrowth. **A)** Cell cycle distribution of AUTS2-deficient human neural progenitor cells (hNPCs). The figure likely shows flow cytometry results, demonstrating the percentage of cells in different cell cycle phases (G1, S, G2/M). n=3 cell lines in triplicate. *p<0.05; **p<0.01; ***p<0.001; ****p<0.0001. **B)** Quantitative comparison of neurosphere diameters 15 days in culture, indicating potential self-renewal or proliferation capacity alterations. p-values were determined by unpaired t-test with Welch’s correction (n=3 cell lines, 8 replicates). *p<0.05; **p<0.01; ***p<0.001; ****p<0.0001. **C)** Wound scratch assay conducted using the Incucyte® system. p-values were determined by unpaired t-test with Welch’s correction (n=3 cell lines, 8 or 10 replicates). *p<0.05; **p<0.01; ***p<0.001; ****p<0.0001. **D)** Neurite outgrowth from NPCs after 1 week in neuronal differentiation media. The figure compares the average length of neurites. p-values were determined by an unpaired t-test with Welch’s correction (n=3, 15 replicates). Progenitor cells to mature neurons. The diagram highlights key morphological and molecular changes occurring at each stage, including cell fate specification, neurite outgrowth, and synapse formation. Red: Cell membrane stain (Neurite Outgrowth Staining Kit, Thermofisher), Magenta: ImageJ Neurite Tracing. *p<0.05; **p<0.01; ***p<0.001; ****p<0.0001.

### WNT7A treatment rescues the cellular phenotype caused by AUTS2 deficiency

Integration of ChIP, eCLIP, and RNA-seq datasets revealed a significant enrichment of genes involved in the Wnt signaling pathway (**Fig. 5E**). RNA-seq analysis identified differentially expressed genes in AUTS2 knockdown NPCs associated with both canonical (β-catenin-dependent) and non-canonical (β-catenin-independent) Wnt signaling, which were mapped onto a comprehensive pathway schematic and STRING analysis (**Fig. 7A, B**). These visualizations highlighted widespread downregulation of Wnt pathway components (**Supplementary Fig.5 and Supplementary Table 5)** with 74 out of 89 genes downregulated, including WNT ligands (examples: WNT1, 7A, 7B, 9A, 9B, 10B), frizzled receptors (examples: FZD1–4, 6, 7, 9, 10), and both VANGL family members (VANGL1 and VANGL2) in AUTS2-deficient NPCs (**Fig. 7A and Supplementary Fig.5**). These molecular disruptions likely contribute to the observed impairments in cell proliferation, migration, and neurite outgrowth observed in AUTS2 knockdown NPCs (**Fig. 6**). Among the affected genes, *WNT7A* was the most significantly downregulated gene in AUTS2 knockdown NPCs and showed direct regulation by AUTS2 at both the DNA and RNA levels (**Figs. 5D, F**). To explore the functional relationship between AUTS2 and WNT7A, and to assess their role in neurogenesis, we performed rescue experiments using recombinant human WNT7A protein. Treatment with WNT7A restored impaired cell migration in neurosphere-derived NPCs and enhanced neurite outgrowth and arborization in AUTS2-deficient neurons (**Figs. 7C, D**).

**Fig. 7:**
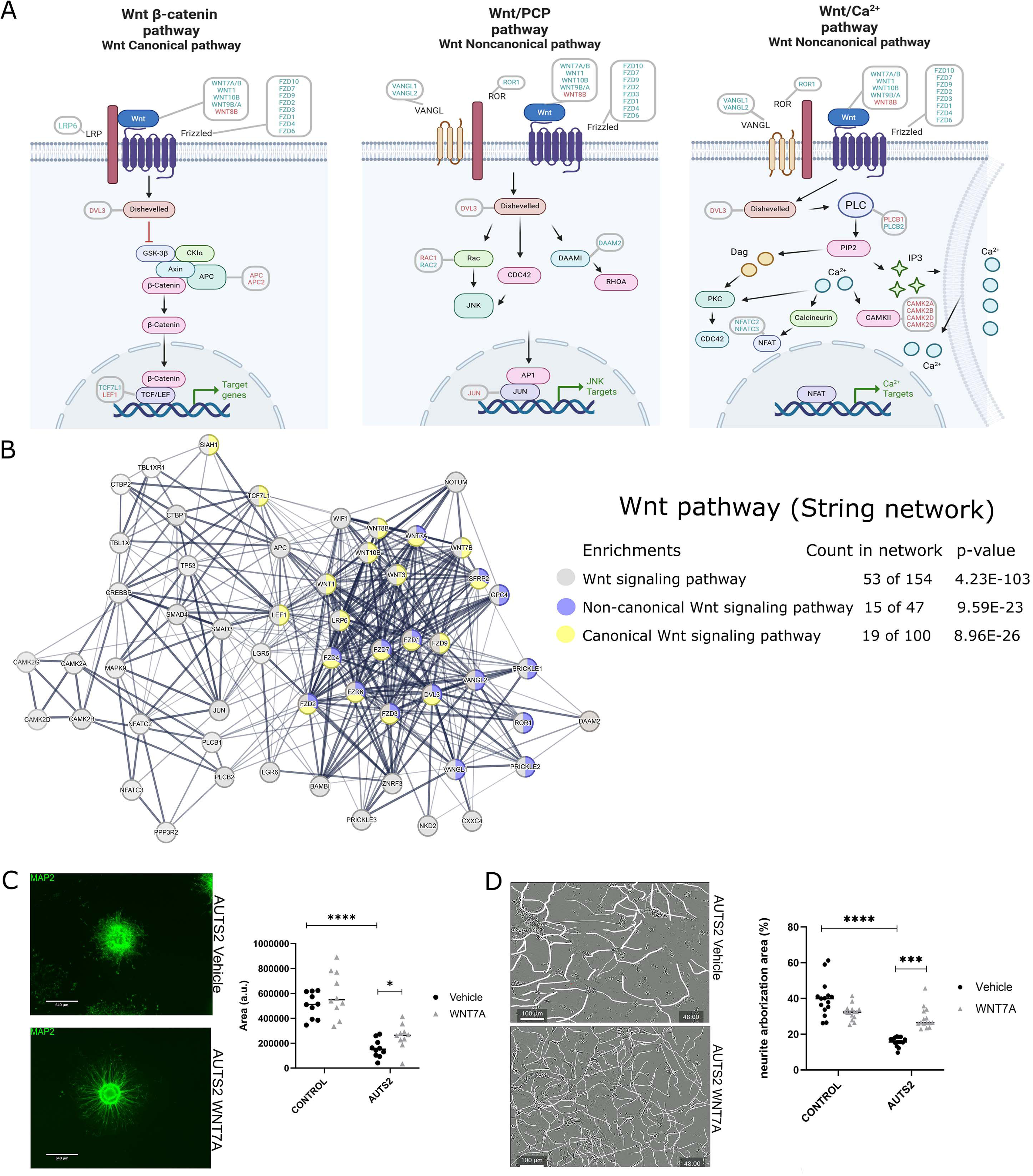
WNT7A treatment rescues the cellular phenotype caused by AUTS2 deficiency. **A)** Schematic representation of the canonical and non-canonical Wnt signaling pathways and its major branches. i) Wnt/β-catenin canonical pathway: This pathway is characterized by the stabilization of β-catenin, which then translocates to the nucleus to regulate gene expression; ii) Wnt/PCP non-canonical pathway: This pathway regulates cell polarity and cytoskeleton organization independently of β-catenin; iii) Wnt/Ca^2+^ non-canonical pathway: This pathway involves the release of intracellular calcium and activation of calcium-dependent signaling molecules. Proteins highlighted in red indicate upregulation, while those in blue indicate downregulation in AUTS2 knockdown NPCs. **B)** STRING analysis of the gene-gene interaction network for Wnt signaling components. This network, derived from RNA-seq analysis of AUTS2 knockdown NPCs versus controls, demonstrates the involvement of both canonical and non-canonical Wnt pathways, with p-values of 8.96e-26 and 9.56e-23, respectively. Gray nodes represent general Wnt signaling components, while yellow indicates the canonical pathway and purple the non-canonical pathway. **C)** Representative images and total area quantification of neurospheres showing outgrowth migration of NPCs with AUTS2 knockdown. p-values were determined by an unpaired t-test with Welch’s correction (n=3). *p<0.05; **p<0.01; ***p<0.001; ****p<0.0001. **D)** Figure illustrates the neurite arborization patterns of NPCs with AUTS2 knockdown. The figure includes representative images showing the complexity of the neuronal branching and a quantitative graph summarizing the data. The results were statistically analyzed using an unpaired t-test with Welch’s correction on three independent biological replicates (n=3). *p<0.05; **p<0.01; ***p<0.001; ****p<0.0001.

Collectively, these findings demonstrate that AUTS2 regulates neurodevelopmental processes in part through modulation of Wnt signaling. Furthermore, identifying genes co-regulated by AUTS2 at both transcriptional and post-transcriptional levels (such as *WNT7A*) may offer promising targets for functional rescue strategies.

## Discussion

AUTS2 is an unusual neurodevelopmental regulator with diverse molecular functions that appear to shift dynamically as brain development progresses, depending on both the cellular context and developmental stage. Previous studies in mouse and human models pointed to a particularly important function in neural progenitor cell proliferation, migration and neuronal differentiation ^6,8,10,16–18^. In this study, we present the first genome-wide mapping of AUTS2’s DNA and RNA binding landscape in human cortical NPCs, showing that AUTS2 dual binding regulates networks of genes in neurodevelopment, neurodevelopmental disease and Wnt signaling pathway. Knockdown of AUTS2 led to widespread changes in gene expression, predominantly affecting processes related to nervous system development, DNA methylation, and RNA metabolism, in agreement with AUTS2’s chromatin and RNA interactions. Our integrative analysis identified a set of 39 genes that were dysregulated at both transcriptional and post-transcriptional levels, and enriched for functions in neuronal differentiation and Wnt signaling; among these, *WNT7A* emerged as the most strongly repressed upon AUTS2 knockdown. AUTS2 deficiency resulted in impaired proliferation and differentiation of neural progenitors, defects that were rescued by WNT7A supplementation. These results advance our understanding of the mechanisms by which AUTS2 coordinates human cortical development and identify WNT7A from the Wnt pathway as a potential target for addressing AUTS2-related deficits.

We used genome-wide eCLIP profiling to identify direct RNA binding targets of AUTS2 in human cortical NPCs and to provide new insight into its role in mRNA processing. AUTS2 was found to bind preferentially near the 5’ and 3’ untranslated regions and to intronic regions. This multi-site binding pattern is uncommon because most RBPs localize to only one or two transcript regions ^30^. However, such a distribution resembles that of RBPs functioning co-transcriptionally to regulate mRNA processing as transcription proceeds. Examples of proteins with similar patterns include the histone modifier zinc finger proteins ZNF473 and PHF7 ^21^, ALYREF from the TREX complex which is involved in RNA export ^31^, and the splicing regulator SRSF7 ^30^. The ability of AUTS2 to function in synchronizing co-transcriptional RNA processing is in line with its established role in chromatin modification, and is ultimately a necessity for proper neurodevelopmental regulation. Consistent with these findings, we observed that AUTS2 knockdown NPCs display significant downregulation of genes linked to RNA metabolism and mRNA processing. On this account, our results confirm and expand the study of Castanza et al. ^9^ in the rodent model, which showed AUTS2’s association with RNA-protein complexes and regulation of transcripts essential for neurogenesis and splicing in mouse cortex. Therefore, our work broadens the molecular repertoire of AUTS2 functions, placing it among neurodevelopmentally relevant RBPs such as MECP2, FMRP, and SRRM4, whose dysfunction is strongly linked to intellectual disability and ASD ^32^. The addition of AUTS2 to this group underscores the importance of post-transcriptional regulation in cortical development and suggests that disruption of AUTS2’s RNA-binding activity may contribute to neurodevelopmental pathology by altering target RNA networks.

In contrast to non-human models like mouse and zebrafish, previous studies using human cortical organoids demonstrated perturbations in Wnt/β-catenin signaling in AUTS2-deficient models.^11,17,18,33^ This points to a potentially human-specific regulatory function for AUTS2 in Wnt pathway modulation. However, these human organoid studies revealed distinct context-dependent consequences: global hyperactivation of canonical Wnt/β-catenin signaling upon AUTS2 loss, while the T534P missense mutation specifically downregulated Wnt pathway transcription in neural progenitors as shown by single-cell sequencing.^17^ ^18^ In our study, we further identified downregulation of five genes involved in Wnt signaling - *WNT7A, HES5, ZNF703, VANGL2* and *SOX11* - in AUTS2-deficient NPCs that are also AUTS2 targets at DNA and RNA levels, each with established roles in neurogenesis, neuronal differentiation, and migration. HES5 facilitates neuronal differentiation from human NPCs through WNT3a-dependent regulation of non-canonical Wnt and NOTCH pathways ^34^. ZNF703 and SOX11 also promote neurogenesis in progenitor cells via transcriptional mechanisms ^35–37^. VANGL2 functions as a core component of the non-canonical Wnt/planar cell polarity (PCP) pathway, acting downstream of WNT7A and is essential for neural development and migration ^38^.

Mechanistically, our data demonstrate direct transcriptional and post-transcriptional control of multiple Wnt pathway genes by AUTS2 and show that WNT7A supplementation can rescue the proliferation, migration, and neurite outgrowth deficits observed in AUTS2-deficient human NPCs. An open question remains: does AUTS2-driven regulation of WNT7A control canonical Wnt/β-catenin signaling, Wnt/PCP, or both? Given that VANGL1 and VANGL2 are also repressed and PCP signaling underlies neurodevelopmental processes such as migration and axon guidance, categories enriched among AUTS2 DNA and RNA targets, our findings support a model in which AUTS2 coordinates both branches of Wnt signaling in human cortical NPCs.

A major strength of our study is its integrated analysis of AUTS2’s activity at chromatin, RNA, and transcriptional levels, revealing convergent regulation of Wnt signaling pathways in human NPCs. This was possible because we took advantage of a simple and controlled system, monolayer cultures of neural progenitor cells, which allowed us to focus precisely on AUTS2’s regulatory role during an early and critical phase of neurogenesis. By restricting analysis to NPC lines, we minimized confounding factors present in more complex organoid systems and directly tracked cell proliferation, migration, and neurite outgrowth. Such early events are essential for the proper establishment of the cortical layers, as the balance between progenitor proliferation and neuronal migration determines both neuronal diversity and laminar organization ^39,40^. Defective regulation at this stage often results in cortical malformations, epilepsy, microcephaly, and neurodevelopmental disorders like intellectual disability and ASD ^41–44^.

AUTS2 is found in protein complexes associated with chromatin and RNA and it’s striking that other genes that interact with or regulate AUTS2 have also been implicated in neurodevelopmental disorders. For example, mutations affecting the catalytic and regulatory subunits of kinase CK2 cause two distinct neurodevelopmental syndromes ^45^; CK2 is part of the polycomb complex PRC1 that modifies chromatin associated with AUTS2. The transcription factor TBR1 that regulates AUTS2 plays a pivotal role in cortical development, and *TBR1* mutations are linked to intellectual disability and ASD ^46^. Similarly, pathogenic mutations in *MED13L* gene cause a syndrome with intellectual disability and the protein primes the transcriptional activation of genes involved in neurogenesis in the developing cortex including AUTS2 ^47^. The convergence of diverse neurodevelopmental risk genes on pathways controlled by AUTS2 highlights the importance of understanding the dynamic molecular interactions and regulatory mechanisms underlying early brain development. Collectively, these findings further reinforce that disruption of AUTS2-centered networks can contribute to a broad spectrum of neurodevelopmental disorders and suggest that therapeutic strategies targeting such pathways may benefit individuals with related conditions.

## Methods

### hESC cell culture and NPC differentiation

hESC H9 line (WA09; WiCell Research Institute, Inc.; CVCL_9773) and AUTS2 KD-derivatives were maintained in mTeSR Plus medium (STEMCELL Technologies, #100-0274) on Cultrex Reduced Growth Factor Basement Membrane Extract (R&D Systems, Catalog #: 3433-010-01)-coated plates, with every other day medium changes. Cells were passed at 80-90% confluency using Gentle Cell Dissociation Reagent (STEMCELL Technologies, #100-0485). Cell lines were screened monthly for Mycoplasma contamination using MycoAlertTM PLUS Mycoplasma Detection Kit capable and consistently tested negative (Lonza, Catalog #: LT07-710).

Pan cortical NPCs were generated according to published protocols ^41,48^. Briefly, embryonic bodies (EBs) were formed from hESC colonies detached using Collagenase IV (Thermo Fisher Scientific, Catalog number 17104019), resuspended in mTeSR Plus with 10 mM Rock inhibitor (Y-27632 dihydrochloride, Tocris Bioscience, #1254), and plated onto low-adherence dishes. The next day, media was changed to N2B27 media (Dulbecco’s modified Eagle’s medium/F12 Glutamax medium, Thermo Fisher Scientific #10565018) supplemented with 1x N2 supplement (Thermo Fisher Scientific #17502048) and 1x B27 supplement (Thermo Fisher Scientific #17504044) supplemented with 500 ng/ml human noggin (R&D Systems, Catalog #: 6057-NG). After 10 days, the EBs were plated onto 10 μg/ml poly-ornithine (Sigma Aldrich, #27378-49-0)/ 5 μg/mL laminin (Invitrogen, #23017-015)-coated dishes in the same media. Rosettes were manually selected after 7 days, dissociated with accutase (STEMCELL AT-104) and plated into new poly-ornithine (PLO)/laminin-coated plates in N2B27 with 20 ng/ml FGF2 (R&D Systems, Catalog #: 223-FB) and 1 μg/ml laminin. Homogeneous populations of NPCs were obtained after 1-2 passages with Accutase in the same media.

### CRISPR/Cas 9 genome editing for *AUTS2* knockdown

Two sgRNAs targeting the *AUTS2* gene were designed for high efficiency and minimal off-target effects using Benchling (www.benchling.com/crispr) (**Table 1**) and cloned into a PX459 Cas9-expressing plasmid (https://www.addgene.org/62988/). 1.5-2 million H9 ESCs were electroporated with 4-5 μg of PX459-sgRNA plasmid using the Cell Line Nucleofector® Kit V (Lonza, Catalog #: VCA-1003) in the Nucleofector™ System II (Lonza), then the cells were plated with 10 mM Rock inhibitor (Y-27632 dihydrochloride, Tocris Bioscience, #1254). After 24 h, cells were treated with 0.25 μg/ml puromycin for 1-2 days to eliminate non-transfected cells. Puromycin-resistant cells were dissociated using Gentle Cell Dissociation Reagent (STEMCELL Technologies, #100-0485). Clonal picking relies on seeding cells at a very low density (3-5 cells/ml) to facilitate the isolation of individual cells. A 100 µl aliquot of this diluted cell suspension was then plated into 96-well plates. Single cells within the wells were identified and marked as positive using a microscope. Following Cas9-mediated cleavage and successful homologous recombination repair, hESC clones containing targeted insertions/deletions (INDELs) within the AUTS2 gene were isolated and expanded for further analysis. We obtained three AUTS2 gene clones with mutations that resulted in premature stop codons (Fig. 3C): clone 3.1 (AUTS2 KD1) presented an INDEL in exon 9, with one allele containing a 1 base pair (bp) insertion and the other a 13 bp deletion; the other two clones, both with INDELs in exon 12, were clone 5.2 (AUTS2 KD2), which had one allele with a 2 bp insertion and the other with a 2 bp deletion, and clone 7.2 (AUTS2 KD3), which contained an 8 bp insertion in one allele and a 16 bp deletion in the other allele. The control lines (Control C1, C2, and C3) used in this study were generated using the same cloning process, but with non-targeting control gRNA (scramble RNA) sequences that do not match any sequence in the human genome.

**Table 1.**
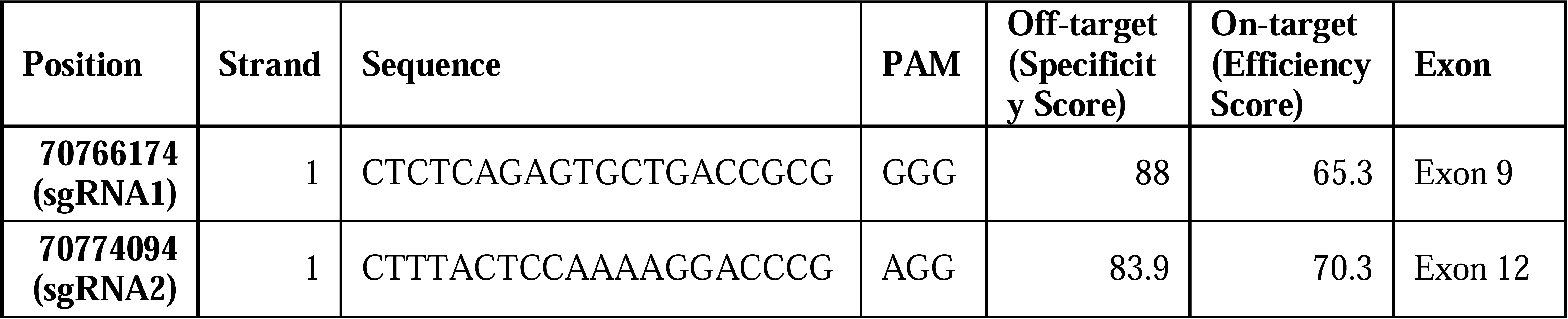
Single guide RNAs for *AUTS2* knockdown expression.

We screened sgRNA1 and sgRNA2 clones for CRISPR/Cas9-induced INDELs using the NGS amplicon-EZ platform. Briefly, we extracted genomic DNA and performed a PCR to amplify the 270bp (sgRNA1) and 320 bp region (sgRNA2) targeted by the sgRNAs (**Table 2**). Partial Illumina adaptors (Forward: 5’-ACACTCTTTCCCTACACGACGCTCTTCCGATCT-3’; Reverse: 5’-GACTGGAGTTCAGACGTGTGCTCTTCCGATCT-3’) were used for paired-end sequencing with a 2×250 bp configuration, generating approximately 50,000 reads per sample. We then aligned these sequences to the sgRNA1 and sgRNA2 reference sequences using Geneious Prime software to identify INDELs. An *in silico* analysis determined the effect of these mutations on the protein, and we selected those clones for further analysis with any mutations that resulted in a frameshift and subsequent protein truncation.

**Table 2.**
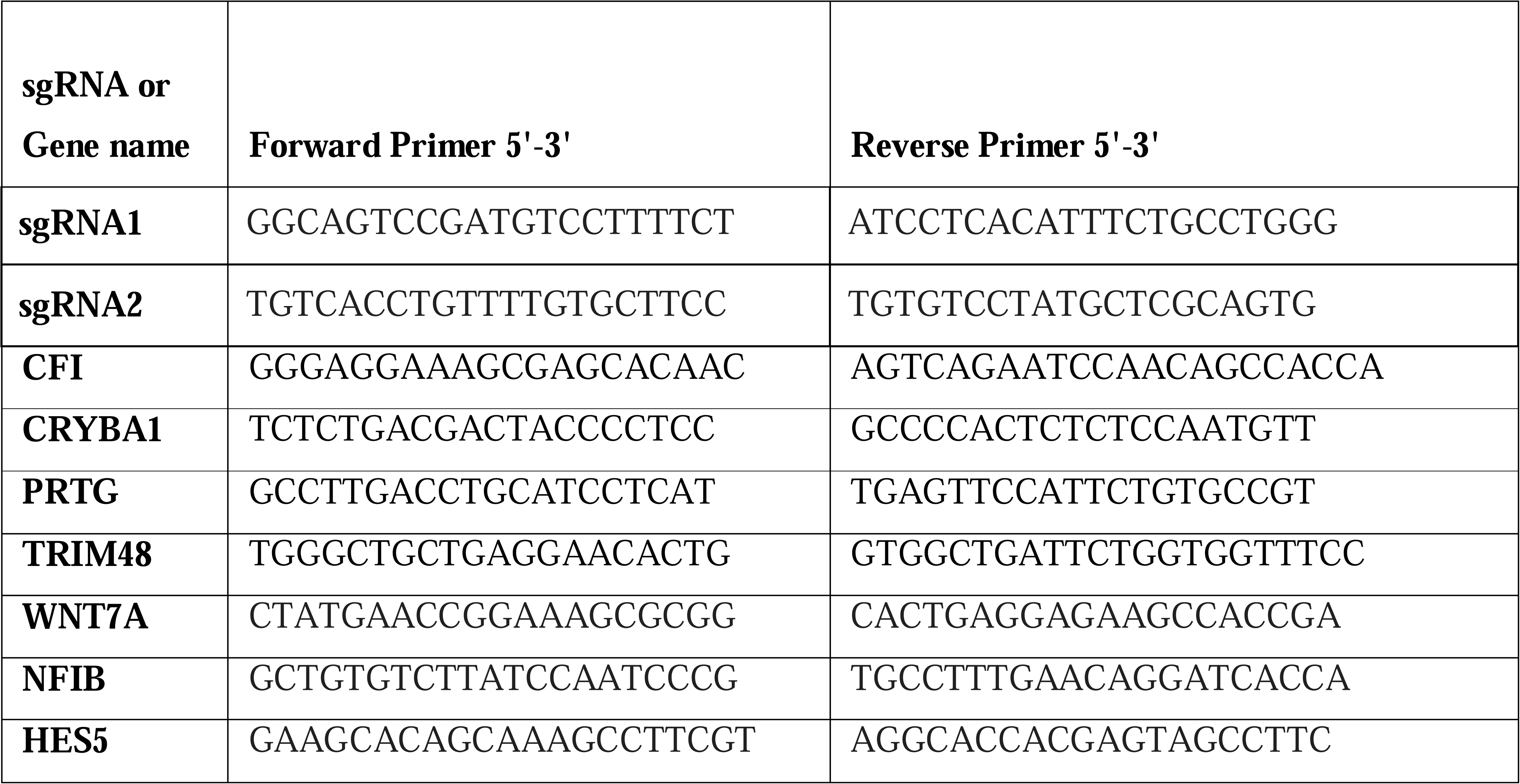
Primers used in this study.

### RT-PCR

NPCs were lysed in Trizol (Thermo Fisher Scientific #15596026) and total RNA was extracted by using RNA Clean & Concentrator-5 (Zymo, #r1014). A reverse transcriptase reaction was carried out using the SuperScript III First-Strand Synthesis System for the RT-PCR kit (Invitrogen, # 18080-051) using the primers listed on **Table 2**. To test the expression, quantitative PCR was performed by using SYBRGreen Universal Master Mix (Applied Biosystems, # 4309155) on QuantStudio 5 qPCR machine from Applied Biosystems. Each cell line was run with two biological replicates and three technical replicates each. GraphPad Prism was used to plot the graphs and for statistical analysis.

### Western blotting

To extract total protein, cultured cells from a 6-well plate were washed with PBS and lysed in protease inhibitor-supplemented RIPA buffer (EMD Millipore, #20-188). The lysates were incubated on ice for 20 min and then centrifuged at 14,000 rpm for 20 min at 4°C to clarify the supernatants. Protein concentration was determined in the supernatants using a Quick Start Bradford protein assay (Bio-Rad, #5000205) with a Promega Glomax microplate reader. Proteins were separated by SDS-PAGE in Bolt™ Bis-Tris Plus Mini Protein 4-12% Gels (Invitrogen, #NW04120BOX) with Bolt MOPS SDS running buffer (Life Technologies, #B0001). The proteins were transferred onto PVDF membranes using NuPAGE™ Transfer Buffer (Invitrogen, #NP00061) with 20% methanol. The primary antibodies used for immunoblotting were anti-AUTS2 (Sigma, #HPA000390) and anti-GAPDH (Fitzgerald, #10R-G109a). Primary antibody binding was detected with horseradish peroxidase-conjugated anti-species and ECL luminescence solution (Millipore, #WBKLS0500)). All samples were analyzed in triplicate for accuracy.

### RNA ligase-mediated rapid amplification of cDNA ends (RLM-RACE)

To identify the presence of an alternative TSS at exon 9, we performed the rapid amplification of 5’ cDNA ends (5’ Race) assay. This assay was performed according to the FirstChoice 5’ RLM RACE Assay (Invitrogen, #AM17000) with slight modifications. For nested PCR, *AUTS2* gene-specific primers (Table 3) and Q5 High-Fidelity 2X Master Mix (New England Biolabs, #M0492S) were used. Inner PCR product was run on an agarose gel, followed by band excision and DNA extraction with Zymoclean Gel DNA Recovery Kit (Zymo, #D4007). The sequence was validated by Sanger sequencing. **Table 3** lists the AUTS2 gene-specific primers used for the RACE assay nested PCR, as previously described by Beunders et al. (2013).

**Table 3.**
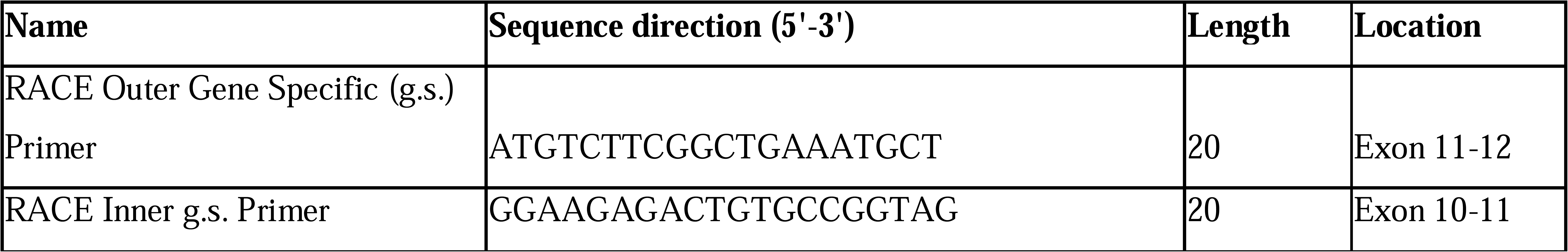
AUTS2 gene-specific primers were used for the RACE assay nested PCR.

### ChIP-Seq

ChIP-sequencing was performed according to a previously established protocol with slight alterations ^49^. NPCs from H9 line were cross-linked with 1% formaldehyde (methanol free) for 10 min, followed by quenching with 2.5 M glycine to final concentration of 125 mM for 5 min at room temperature (RT). Samples were lysed (300 mM NaCl, 10 mM Tris-HCl pH 8.0, 1 mM EDTA pH 8.0, 0.1% Na-deoxycholate, 0.1% SDS, 1% Triton X-100, 0.25% sarkosyl, 1X protease inhibitor cocktail) and sonicated to obtain 200-500 bp chromatin fragments. For each IP reaction, 2.5 μg of anti-AUTS2 antibody (Sigma, # HPA000390) bound to protein A Dynabeads complex (Invitrogen, #10001D) was used. The pull-down chromatin was reverse cross-linked, followed by 0.2 μg/μl RNase A (Epochlifescience, #B408-100) and 0.2 μg/μl proteinase K (Meridian Life Sciences, #C788T38) treatment. The DNA was purified and quality checked on the TapeStation system from Agilent Technologies. Libraries were constructed with the NEBNext Ultra II DNA Library Prep Kit for Illumina (New England Biolabs, NEB #7645L) and sequencing was performed using an Illumina HiSeq 2500 Sequencer with 50 bp single-end reads and a minimum depth of 70 million reads per sample. Raw FASTQ files were aligned to the GRCh38 (hg38) reference genome using STAR aligner 2.7.10 ^50^. Following alignment, narrow AUTS2 binding peaks were identified using the MACS2 peak calling software ^28^. The resulting narrowPeak files were then loaded into R using the DiffBind package 3.14.0 ^51^. Consensus peaks were identified as those present in all 3 of the CHIP replicates performed. Peaks were annotated using ChIPseeker 1.40.0 ^52^ and the org.Hs.eg.db database. GO enrichment analysis was performed on CHIP target genes using clusterProfiler 4.12.6 ^53^. Enriched AUTS2 DNA binding motifs were identified by HOMER 4.11 ^19^. Additional analyses and the generation of plots were performed using custom R scripts.

### eCLIP

Neural progenitor cells (NPCs were cultured in 15□cm dishes to ∼90% confluency and crosslinked with 400□mJ/cm² UV-C light. Cells were lysed in buffer (50□mM HEPES pH□7.4, 150□mM NaCl, 0.5% NP-40, 0.1% SDS, 0.5□mM DTT, 1□mM PMSF, and protease inhibitors). Lysates were centrifuged at 10000xg and the supernatant was used for immunoprecipitation with anti-AUTS2 antibody (Sigma, #HPA000390) pre-coupled to Dynabeads (Thermo Fisher). Incubation was performed overnight at 4□°C. Two percent of each sample was saved as input. RNA-protein complexes were washed sequentially with high-salt buffer (50□mM HEPES pH□7.4, 1□M NaCl, 1□mM EDTA, 0.1% SDS, 1% NP-40), low-salt buffer (same as above with 150□mM NaCl), and PNK buffer (50□mM Tris pH□7.4, 10□mM MgCl□, 0.5% NP-40). Both IP and input RNA were dephosphorylated using FastAP (Thermo Fisher, #EF0651) and T4 PNK (New England Biolabs, #M0201S), then ligated to a 3′ RNA adaptor with T4 RNA ligase (New England Biolabs, #M0204S). To confirm IP efficiency, 10% of each IP and input was analyzed by Western blot. The remaining 90% was resolved by PAGE, transferred to nitrocellulose, and the protein–RNA complex region excised. Proteinase K digestion released RNA, which was purified. Input RNA was processed in parallel. Reverse transcription was performed using AffinityScript (Agilent, #200436), followed by ExoSAP-IT (Affymetrix, #78201.1.ML) treatment. RNA was degraded by alkaline hydrolysis, a 3′ DNA adaptor was ligated, and libraries were amplified by PCR, gel size-selected, and sequenced on an Illumina HiSeq platform. Each experiment included IP from two independent biological replicates with matched input controls.

Raw FASTQ files were processed and analyzed using Skipper 1.99.0 ^29^, an end to end analysis pipeline for eCLIP data. This pipeline includes alignment to the GRCh38 (hg38) reference genome via STAR 2.7.10 ^50^, the identification and annotation of AUTS2 binding peaks via Skipper’s core peak calling method, and the identification of consensus peaks (peaks found in both eCLIP replicates). Enriched AUTS2 RNA binding motifs were identified by HOMER 4.11 ^19^. GO enrichment analysis was performed on eCLIP target genes using clusterProfiler 4.12.6 ^53^. Additional analyses and the generation of plots were performed using custom R scripts.

### RNA sequencing

Total RNA was isolated from 3-5 million NPCs using TRIzolReagent (Invitrogen), and purified using the RNA Clean & Concentrator kit (Zymo Research, cat #r1014) following the manufacturer’s protocol. RNA was quantified using a Qubit® 3.0 Fluorometer (Thermo Fisher Scientific) and quality control was performed in Tapestation. Libraries from samples with ≥9 RIN value were prepared using Truseq’s Stranded mRNA(PolyA+) Library Prep kit (Catalog #20020595) and sequenced on a NextSeq_HighOutput machine. The NPC samples were sequenced with 75bp long paired-end reads with a minimum depth of 57 million reads per sample. All samples were sequenced in duplicate. Raw FASTQ files were aligned to the GRCh38 (hg38) reference genome using STAR aligner 2.7.10 ^50^. Read counts for each feature/gene were then quantified using featureCounts ^54^, and genes with significant differential expression were then identified using DESeq2 1.44.0 ^55^. GO enrichment analysis was performed on differentially expressed genes using clusterProfiler 4.12.6 ^53^. Additional analyses and the generation of plots were performed using custom R scripts.

### Immunofluorescence

NPCs were seeded onto Lâmina EZ Millicell® (Merck Millipore, #PEZGS0816), designed for microscopy, and allowed to reach 60-70% confluence. Subsequently, cells were fixed with 4% paraformaldehyde for 20 min at RT, followed by 3 washes with PBS. Cells were permeabilized with 0.25% Triton X-100 for 10 min and then were blocked with 1% horse serum for 1 h at RT. Overnight incubation was done at 4°C with Primary antibodies: mouse anti-Nestin (1:500, EMD Millipore MAB5326) and rat anti-SOX2 (1:750, Invitrogen 14-9811-82). Following washes, cells were incubated with secondary antibodies in the dark for 1 h at RT. Secondary antibodies: donkey anti-rat AlexaFluor 488 and donkey anti-mouse AlexaFluor 647 (Invitrogen, 1:200) were used. Finally, nuclei were counterstained with DAPI for 10 min before visualization using a fluorescence microscope (ECHO).

### Cell migration and invasion assay

NPCs were seeded on PLO/laminin-coated 96-well ImageLock tissue culture plates (Sartorius, #4379), and after 24h, a uniform scratch wound was created in each well using a WoundMaker (IncuCyte® WoundMaker). Following two washes with culture medium to remove detached cells, fresh medium was added (N2B27 supplemented with 20 ng/ml FGF2). Cell migration into the wound area was then monitored by time-lapse imaging using either IncuCyte or CELLCYTE X systems. Images were captured every 1-2 h for up to 72 h, typically using a 10X objective (wide mode). Each experimental condition consisted of 8 or 10 replicates (n=8 or n=10). Collected images were analyzed to quantify wound closure using ImageJ software ^56,57^.

### Cell cycle

NPCs were used at ∼85% confluency and were dissociated into single cells by using accutase. Following ethanol fixation for 2 h on ice, cells were incubated overnight at 4°C with propidium iodide (50 µg/mL, Invitrogen, Cat# P1304MP). Fluorescence data were subsequently acquired via flow cytometry analysis using a BD Influx™ Cell Sorter, performed by the Flow Cytometry Core Facility of the Salk Institute.

### Generation of neurospheres

Neural Progenitor Cells (NPCs) were gently separated into a single-cell suspension using Accutase (Thermo Fisher Scientific, MA, USA). This suspension was used for neurosphere generation, which involved culturing the cells for 72 hours in NPC medium on an orbital shaker continuously running at 90 rpm to promote self-aggregation. Following this initial aggregation period, the resulting neurospheres underwent a 3-day maturation phase in adherent conditions and static incubator, maintained at 37°C with 5% CO2 and NPC medium. Adherent neurospheres were fixed on day 3 and immunofluorescence was performed to detect neurite extension.

### Neurite length

One week differentiated neurons (N2B27 with the addition of 1 μg/ml laminin, 20 ng/ml BDNF, 20 ng/ml GDNF, and 500 μg/ml cyclic AMP) were plated as single cells on a 24-well plate coated with PLO/laminin and stained according to the Neurite Outgrowth Staining Kit (Life Technologies, #A15001). Images were taken at 10X objective on Echo microscope, and for each line, 15 different neurites were used for analysis using ImageJ software.

### Statistics and reproducibility

Our findings are reported as mean values, accompanied by either the standard deviation (SD) or the standard error of the mean (SEM). The criterion for statistical significance was a P value of less than 0.05. The specific P values and the statistical tests employed for each experiment are detailed in the corresponding figure legends. These analyses—including two-tailed one-sample tests, unpaired two-tailed Student’s t-tests, and two-way ANOVA followed by post hoc tests—were all conducted using GraphPad Prism v9.0. The number of biological replicates (n values) for each experiment is also provided in the figure legends.

## Data availability

All raw and processed eCLIP, CHIP-seq, and RNA-seq data from this study have been deposited in the Gene Expression Omnibus (GEO). The accession numbers are as follows: GSE308353 for CHIP-seq, GSE304933 for eCLIP and GSE308696 for RNA-seq. Previously published NPC ATAC-seq and histone modification ChIP-seq data were from GSE158382 ^58^.

## Acknowledgments

This work was supported by the Next Generation Sequencing (NGS) Core Facility of the Salk Institute (RRID: SCR_014846 and SCR_026396) with funding from NIH-NCI CCSG P30 CA014195, NIH-NIA San Diego Nathan Shock Center P30 AG068635, the Chapman Foundation, and the Helmsley Charitable Trust. Core personnel include Dr. Elsa Molina, Ling Ouyang, Cristian Quintero, Tzu-Wen Wang, Nina Tonnu, and James Nguyen. ChIP sequencing (Illumina HiSeq platform) was conducted at the IGM Genomics Center, University of California, San Diego, La Jolla, CA. M.C.M. was funded in part by a grant from the California Institute for Regenerative Medicine (DISC4-16295). The contents of this publication are solely the responsibility of the authors and do not necessarily represent the official views of CIRM or any other agency of the State of California.

## Author contributions

V.M.S.C. and M.C.M. conceived the study. V.M.S.C., J.T.W. and L.R.M. designed and conducted the CRISPR/Cas9 experiments. V.M.S.C., M.C.M, A.S., SB., D.X., A.P.D.M., J.T.W., and R.O. carried out cell culture experiments. A.S. and A.C. performed Western blot experiments. A.S. conducted ChIP experiments. S.O., K.F., and G.Y. performed eCLIP experiments and analysis. A.S. performed RACE assays and PCR/qPCR experiments. K.F., V.M.S.C., C.B., G.R.O., S.S., and A.S., conducted bioinformatics analysis. V.M.S.C., A.S., M.J., and J.T.W. analyzed the overall data. V.M.S.C., R.S., and M.C.M. wrote the manuscript. All authors reviewed and approved the final manuscript.

## Supplementary Information

**Supplementary Fig. 1: AUTS2 predominantly binds active chromatin at both promoter and promoter-distal sites. A)** Enrichment of H3K27ac (all active sites) and H3K4me3 (promoters) chromatin marks at AUTS2 peaks. (log2 IP/input, +/-1 kb from peak center). **B)** Enrichment of H3K27ac (active) and H3K27me3 (inactive) chromatin marks at AUTS2 peaks (log2 IP/input, +/-1 kb from peak center).

**Supplementary Fig. 2: Sequence analysis of the *AUTS2* locus in CRISPR/Cas9-edited hESCs cell lines used in this study, showing the identified mutations.**

**Supplementary Fig. 3: 5’ RACE demonstrates the presence of an alternative transcription start site (TSS) in the *AUTS2* gene generating a shorter C-terminal isoform AUTS2-C of approximately 79 kDa, detected by western blot (711 amino acids) in Fig. 3C**.

**Supplementary Fig. 4: Dual DNA/RNA binding by AUTS2 regulates genes involved in neurodevelopmental disorders. A)** Venn diagram illustrating the shared genes between the AUTS2 CLIP-seq and ChIP-seq gene set and three distinct disease-related gene cohorts. These cohorts include genes from the SFARI, ID, and DDD databases. **B)** Gene list displays the gene names from the overlapping set of genes identified in AUTS2 CLIP-seq/ChIP-seq experiments shown in A. **C)** UCSC genome browser visualization of *NSD2* gene locus showing chromatin marks associated with AUTS2 ChIP and eClip peaks. UCSC Genome Browser view of the NSD2 locus (hg38) in neural progenitor cells (NPCs), shown in two genomic windows (chr4:1,863,500–1,865,500 and chr4:1,955,000–1,957,000) for optimal visualization of both eCLIP and ChIP peaks. The RefSeq annotation track shows exons as blue boxes and introns as lines. ChIP-seq peaks are called by MACS2 (see Methods) and appear as black boxes, with normalized coverage (IP in red, input in purple). eCLIP peaks called by Skipper (see Methods) are marked with black boxes, and normalized coverage (IP in red, input in purple). Also shown are ATAC-seq, H3K27ac, H3K27me3 and H3K4me3 tracks. *NSD2* gene encodes for a methyltransferase that has been implicated in modulation of synaptic genes during early brain development ^59,60^ AUTS2 ChIP-seq peaks in the *NSD2* locus were localized within the NSD2 promoter area. ATAC-seq track confirmed that the ChIP peaks coincide with open chromatin, while H3K27ac, H3K27me3, and H3K4me3 tracks illustrate histone modification patterns around the ChIP peak. eCLIP peaks revealed a sharp focal enrichment within exon 15 of *NDS2* gene transcript.

**Supplementary Fig. 5: Differentially regulated genes in AUTS2 KD that participate on the Wnt signaling pathway.** Heatmap displaying the log2 fold change in expression of genes participating in Wnt signaling pathway in AUTS2 KD NPCs compared to control lines (pAdj < 0.05), highlighting widespread downregulation of Wnt pathway components.

## Supplementary Tables

**Supplementary Table 1:** AUTS2 ChIP seq peaks

**Supplementary Table 2:** AUTS2 eCLIP seq peaks

**Supplementary Table 3:** Differentially regulated genes in AUTS2 knockdown after RNA seq experiment.

**Supplementary Table 4:** Intersection between ChIPseq, eCLIP and RNAseq differentially regulated genes in AUTS2 Knockdown.

**Supplementary Table 5:** Differentially regulated genes in AUTS2-defficient NPCs that are involve in Wnt signaling pathway.

## References

1. Sultana, R. et al. Identification of a novel gene on chromosome 7q11.2 interrupted by a translocation breakpoint in a pair of autistic twins. Genomics 80, 129–134 (2002).

2. Oksenberg, N., Stevison, L., Wall, J. D. & Ahituv, N. Function and regulation of AUTS2, a gene implicated in autism and human evolution. PLoS Genet. 9, e1003221 (2013).

3. Kikuchi, Y. et al. Evolutionary constrained genes associated with autism spectrum disorder across 2,054 nonhuman primate genomes. Mol. Autism 16, 5 (2025).

4. Hori, K., Shimaoka, K. & Hoshino, M. AUTS2 gene: Keys to understanding the pathogenesis of neurodevelopmental disorders. Cells 11, 11 (2021).

5. Loberti, L. et al. AUTS2-related syndrome: Insights from a large European cohort. Genet. Med. 27, 101375 (2025).

6. Beunders, G. et al. Exonic deletions in AUTS2 cause a syndromic form of intellectual disability and suggest a critical role for the C terminus. Am. J. Hum. Genet. 92, 210–220 (2013).

7. Sanchez-Jimeno, C. et al. Attention deficit hyperactivity and autism spectrum disorders as the core symptoms of AUTS2 syndrome: Description of five new patients and update of the frequency of manifestations and genotype-phenotype correlation. Genes (Basel) 12, 1360 (2021).

8. Gao, Z. et al. An AUTS2-Polycomb complex activates gene expression in the CNS. Nature 516, 349–354 (2014).

9. Castanza, A. S. et al. AUTS2 regulates RNA metabolism and dentate gyrus development in mice. Cereb. Cortex 31, 4808–4824 (2021).

10. Hori, K. et al. Cytoskeletal regulation by AUTS2 in neuronal migration and neuritogenesis. Cell Rep. 9, 2166–2179 (2014).

11. Jha, U., Kondrychyn, I., Korzh, V. & Thirumalai, V. High behavioral variability mediated by altered neuronal excitability in auts2 mutant zebrafish. eNeuro 8, ENEURO.0493–20.2021 (2021).

12. Monderer-Rothkoff, G. et al. AUTS2 isoforms control neuronal differentiation. Mol. Psychiatry 26, 666–681 (2021).

13. Pang, W. et al. Untangle the multi-facet functions of Auts2 as an entry point to understand neurodevelopmental disorders. Front. Psychiatry 12, 580433 (2021).

14. Shimaoka, K. et al. The microcephaly-associated transcriptional regulator AUTS2 cooperates with Polycomb complex PRC2 to produce upper-layer neurons in mice. EMBO J. 44, 1354–1378 (2025).

15. Liu, S. et al. NRF1 association with AUTS2-Polycomb mediates specific gene activation in the brain. Mol. Cell 81, 4757 (2021).

16. Wang, Q. et al. WDR68 is essential for the transcriptional activation of the PRC1-AUTS2 complex and neuronal differentiation of mouse embryonic stem cells. Stem Cell Res. 33, 206–214 (2018).

17. Fair, S. R. et al. Cerebral organoids containing an AUTS2 missense variant model microcephaly. Brain 146, 387–404 (2023).

18. Geng, Z., Tai, Y. T., Wang, Q. & Gao, Z. AUTS2 disruption causes neuronal differentiation defects in human cerebral organoids through hyperactivation of the WNT/β-catenin pathway. Sci. Rep. 14, 19522 (2024).

19. Heinz, S. et al. Simple combinations of lineage-determining transcription factors prime cis-regulatory elements required for macrophage and B cell identities. Mol. Cell 38, 576– 589 (2010).

20. Van Nostrand, E. L. et al. Robust transcriptome-wide discovery of RNA-binding protein binding sites with enhanced CLIP (eCLIP). Nat. Methods 13, 508–514 (2016).

21. Gosztyla, M. L. et al. Integrated multi-omics analysis of zinc-finger proteins uncovers roles in RNA regulation. Mol. Cell 84, 3826–3842.e8 (2024).

22. Biel, A. et al. AUTS2 syndrome: Molecular mechanisms and model systems. Front. Mol. Neurosci. 15, 858582 (2022).

23. SFARI Gene. Simons Foundation Autism Research Initiative (SFARI) https://gene.sfari.org/database/human-gene/.

24. Deciphering Developmental Disorders (DDD). Sanger institute https://www.sanger.ac.uk/collaboration/deciphering-developmental-disorders-ddd/.

25. Intellectual disability-related genes. Orphanet https://id-genes.orphanet.app/ithaca/.

26. Breast Cancer mutation dataset. Science data bank https://www.scidb.cn/en/detail?dataSetId=d0b2d178e5ce4600b13d9923901c87da.

27. Logan, C. Y. & Nusse, R. The Wnt signaling pathway in development and disease. Annu. Rev. Cell Dev. Biol. 20, 781–810 (2004).

28. Gaspar, J. M. Improved peak-calling with MACS2. bioRxiv (2018) doi:10.1101/496521.

29. Boyle, E. A. et al. Skipper analysis of eCLIP datasets enables sensitive detection of constrained translation factor binding sites. Cell Genom. 3, 100317 (2023).

30. Van Nostrand, E. L. et al. Principles of RNA processing from analysis of enhanced CLIP maps for 150 RNA binding proteins. Genome Biol. 21, 90 (2020).

31. Viphakone, N. et al. Co-transcriptional loading of RNA export factors shapes the human transcriptome. Mol. Cell 75, 310–323.e8 (2019).

32. Li, M. et al. Identification of FMR1-regulated molecular networks in human neurodevelopment. Genome Res. 30, 361–374 (2020).

33. Geng, Z. et al. AUTS2 controls neuronal lineage choice through a novel PRC1-independent complex and BMP inhibition. Stem Cell Rev Rep 19, 531–549 (2023).

34. Mußmann, C., Hübner, R., Trilck, M., Rolfs, A. & Frech, M. J. HES5 is a key mediator of Wnt-3a-induced neuronal differentiation. Stem Cells Dev. 23, 1328–1339 (2014).

35. Da Silva, F. & Niehrs, C. Multimodal Wnt signalling in the mouse neocortex. Cells Dev. 174, 203838 (2023).

36. Kumar, A. et al. Zfp703 is a Wnt/β-catenin feedback suppressor targeting the β-catenin/Tcf1 complex. Mol. Cell. Biol. 36, 1793–1802 (2016).

37. Da Silva, F. et al. Mitotic WNT signalling orchestrates neurogenesis in the developing neocortex. EMBO J. 40, e108041 (2021).

38. Dreyer, C. A., VanderVorst, K. & Carraway, K. L., 3rd. Vangl as a master scaffold for Wnt/planar cell polarity signaling in development and disease. Front. Cell Dev. Biol. 10, 887100 (2022).

39. Jiang, X. & Nardelli, J. Cellular and molecular introduction to brain development. Neurobiol. Dis. 92, 3–17 (2016).

40. Govindan, S. & Jabaudon, D. Coupling progenitor and neuronal diversity in the developing neocortex. FEBS Lett. 591, 3960–3977 (2017).

41. Marchetto, M. C. et al. Altered proliferation and networks in neural cells derived from idiopathic autistic individuals. Mol. Psychiatry 22, 820–835 (2017).

42. Juric-Sekhar, G. & Hevner, R. F. Malformations of cerebral cortex development: Molecules and mechanisms. Annu. Rev. Pathol. 14, 293–318 (2019).

43. Blumcke, I. et al. Neocortical development and epilepsy: insights from focal cortical dysplasia and brain tumours. Lancet Neurol. 20, 943–955 (2021).

44. Mato-Blanco, X. et al. Early developmental origins of cortical disorders modeled in human neural stem cells. Nat. Commun. 16, 6347 (2025).

45. Ballardin, D., Cruz-Gamero, J. M., Bienvenu, T. & Rebholz, H. Comparing two neurodevelopmental disorders linked to CK2: Okur-Chung neurodevelopmental syndrome and Poirier-Bienvenu neurodevelopmental syndrome-two sides of the same coin? Front. Mol. Biosci. 9, 850559 (2022).

46. Chuang, H.-C., Huang, T.-N. & Hsueh, Y.-P. T-brain-1--A potential master regulator in autism spectrum disorders: TBR1 controls expression of autism related genes. Autism Res. 8, 412–426 (2015).

47. Li, J. et al. Regulation of cortical neurogenesis by MED13L via transcriptional priming and its implications for MED13L syndrome. *Commun*. Biol. 8, 1180 (2025).

48. Marchetto, M. C. N. et al. A model for neural development and treatment of Rett syndrome using human induced pluripotent stem cells. Cell 143, 527–539 (2010).

49. Hawkins, R. D. et al. Distinct epigenomic landscapes of pluripotent and lineage-committed human cells. Cell Stem Cell 6, 479–491 (2010).

50. Dobin, A. et al. STAR: ultrafast universal RNA-seq aligner. Bioinformatics 29, 15–21 (2013).

51. Stark, R. & Brown, G. D. DiffBind: Differential binding analysis of ChIP-Seq peak data. (2012).

52. Yu, G., Wang, L.-G. & He, Q.-Y. ChIPseeker: an R/Bioconductor package for ChIP peak annotation, comparison and visualization. Bioinformatics 31, 2382–2383 (2015).

53. Yu, G., Wang, L.-G., Han, Y. & He, Q.-Y. clusterProfiler: an R package for comparing biological themes among gene clusters. OMICS 16, 284–287 (2012).

54. Liao, Y., Smyth, G. K. & Shi, W. featureCounts: an efficient general purpose program for assigning sequence reads to genomic features. Bioinformatics 30, 923–930 (2014).

55. Love, M. I., Huber, W. & Anders, S. Moderated estimation of fold change and dispersion for RNA-seq data with DESeq2. Genome Biol. 15, 550 (2014).

56. Venter, C. & Niesler, C. U. Rapid quantification of cellular proliferation and migration using ImageJ. Biotechniques 66, 99–102 (2019).

57. Cormier, N., Yeo, A., Fiorentino, E. & Paxson, J. Optimization of the wound scratch assay to detect changes in Murine mesenchymal stromal cell migration after damage by soluble cigarette smoke extract. J. Vis. Exp. e53414 (2015).

58. Choi, W.-Y. et al. NEUROD1 intrinsically initiates differentiation of induced pluripotent stem cells into neural progenitor cells. Mol. Cells 43, 1011–1022 (2020).

59. Kinoshita, S. et al. Loss of NSD2 causes dysregulation of synaptic genes and altered H3K36 dimethylation in mice. Front. Genet. 15, 1308234 (2024).

60. Ritchie, F. D. & Lizarraga, S. B. The role of histone methyltransferases in neurocognitive disorders associated with brain size abnormalities. Front. Neurosci. 17, 989109 (2023).

