## Supplementary figures and images for "Human AUTS2 regulates neurodevelopmental pathways via dual DNA/RNA binding"

### Supplemental Figure 1

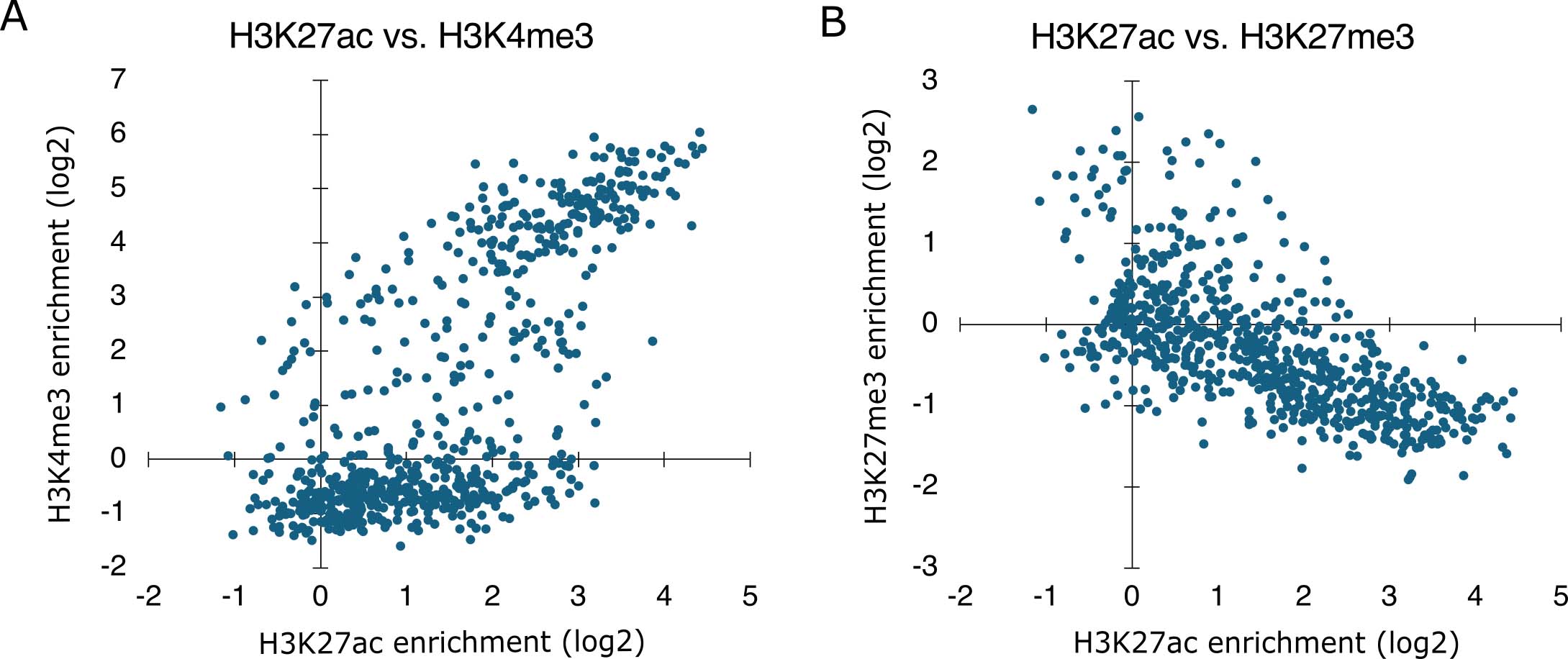

### Supplemental Figure 2

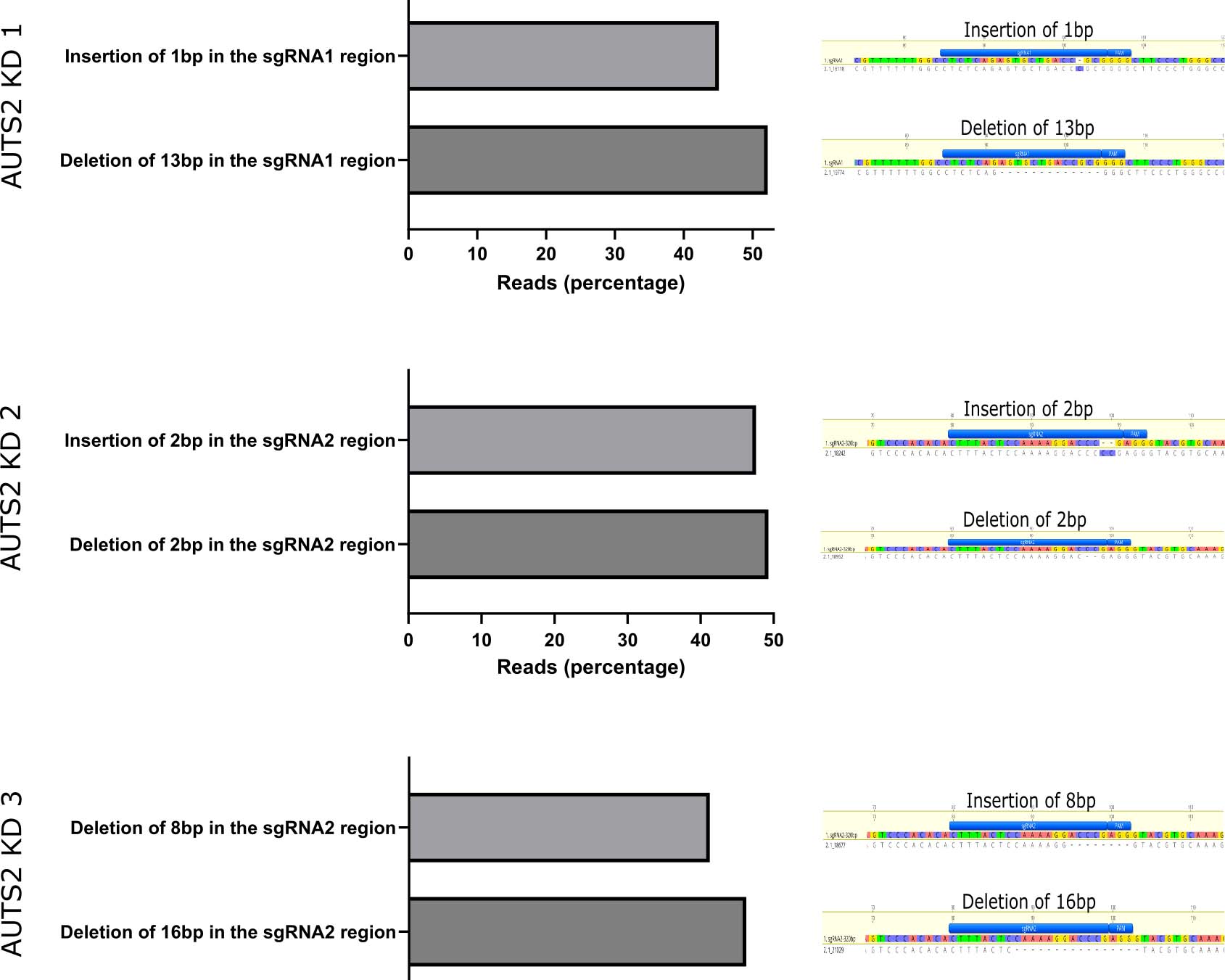

### Supplemental Figure 3

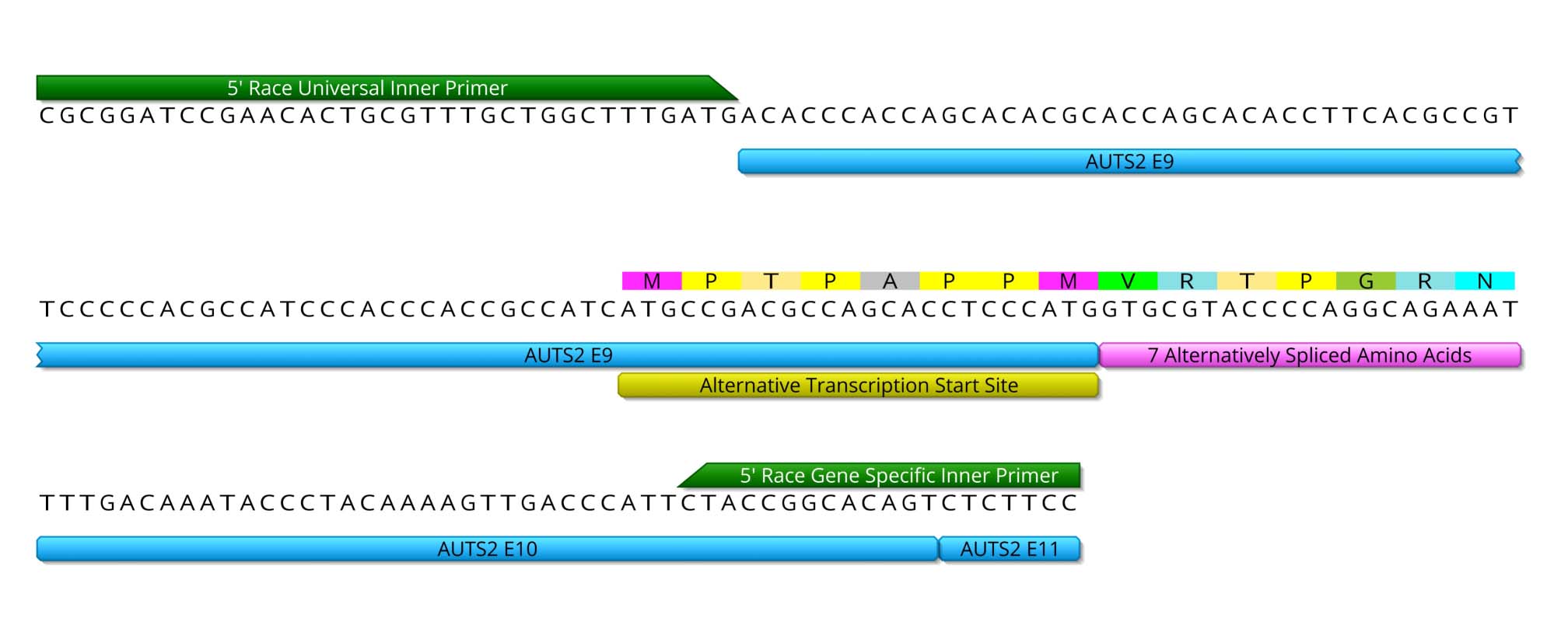

### Supplemental Figure 4

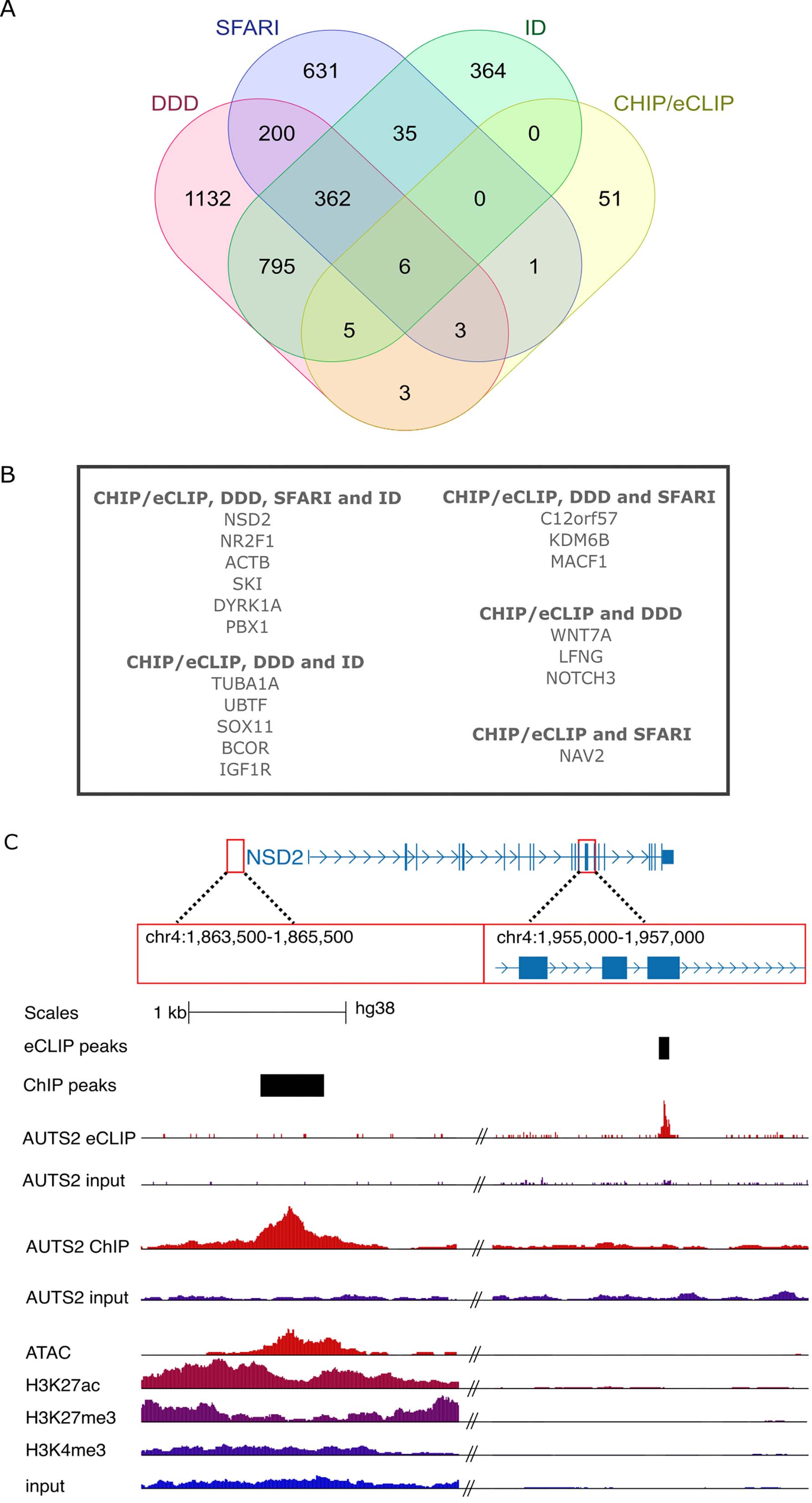

### Supplemental Figure 5

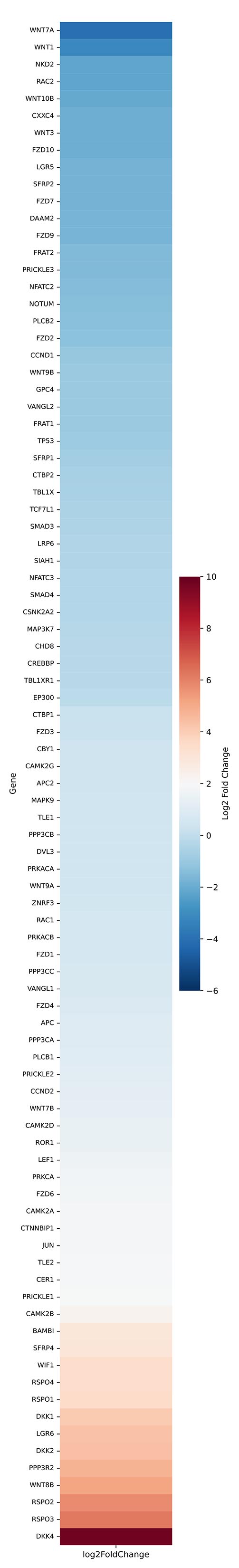
